# SIRV: Spatial inference of RNA velocity at the single-cell resolution

**DOI:** 10.1101/2021.07.26.453774

**Authors:** Tamim Abdelaal, Laurens M. Grossouw, R. Jeroen Pasterkamp, Boudewijn P.F. Lelieveldt, Marcel J.T. Reinders, Ahmed Mahfouz

## Abstract

RNA Velocity allows the inference of cellular differentiation trajectories from single-cell RNA sequencing (scRNA-seq) data. It would be highly interesting to study these differentiation dynamics in the spatial context of tissues. Estimating spatial RNA velocities is, however, limited by the inability to spatially capture spliced and unspliced mRNA molecules in high-resolution spatial transcriptomics. We present SIRV, a method to spatially infer RNA velocities at the single-cell resolution by enriching spatial transcriptomics data with the expression of spliced and unspliced mRNA from reference scRNA-seq data. We used SIRV to infer spatial differentiation trajectories in the developing mouse brain, including the differentiation of midbrain-hindbrain boundary cells and marking the forebrain origin of the cortical hem and diencephalon cells. Our results show that SIRV reveals spatial differentiation patterns not identifiable with scRNA-seq data alone. Additionally, we applied SIRV to mouse organogenesis data and obtained robust spatial differentiation trajectories. Finally, we verified the spatial RNA velocities obtained by SIRV using 10x Visium data of the developing chicken heart and MERFISH data from human osteosarcoma cells. Altogether, SIRV allows the inference of spatial RNA velocities at the single-cell resolution to facilitate studying tissue development.

## 1. Introduction

Single-cell RNA-sequencing (scRNA-seq) enables the study of cellular differentiation dynamics at single-cell resolution^1^. Trajectory inference methods, such as Monocle^2^, DPT^3^ and PAGA^4^, aim to define an ordering of gene expression changes in a certain pool of cells. This ordering potentially reflects a trajectory of the cellular differentiation process. However, scRNA-seq only captures a static snapshot of the cellular states, which represents a major challenge for trajectory inference methods to correctly capture the dynamics of the cellular differentiation process. RNA velocity^5,6^ addresses this challenge by estimating the dynamics of cellular differentiation using the expression balance between unspliced immature and spliced mature mRNA molecules captured by scRNA-seq protocols. Typically, for each cell, the RNA velocity is estimated for each gene individually, which together, predict the future state of the cell. When calculated over all cells, a flow field can be calculated by some averaging of neighboring cells, which can be projected in a low-dimensional visualization space.

To date, the study of cellular differentiation is limited to dissociated cells from scRNA-seq, ignoring the spatial organization of cells in tissues. Taking the spatial context into account can enhance our understanding of cellular differentiation processes^7^. Spatial transcriptomics enable the study of the cellular heterogeneity of complex tissues while retaining spatial information^8,9^. Currently, a wide range of protocols^10–17^ is available which varies in spatial resolution, gene detection sensitivity and number of simultaneously measured genes. In principle, it is possible to apply RNA velocity analysis to spatial transcriptomics measured using sequencing-based protocols, such as 10x Visium and Slide-seq^10,11^, as the spliced and unspliced expression ratios can be directly obtained from the sequencing data^11,18^. However, the spatial resolution of these protocols is currently limited to measuring tissue spots consisting of multiple cells. On the other hand, high resolution protocols, such as seqFISH^16,17^ and HybISS^19^, can provide (sub)cellular resolution but lack the spliced and unspliced counts required to study cellular differentiation using RNA velocity.

We and others have previously shown that high-resolution imaging-based spatial transcriptomics data can be enriched with predicted expression of spatially unmeasured genes through integration with scRNA-seq data measured from the same tissue^20–25^. Building on the same concept, it should be possible to predict the spliced and unspliced expressions of each spatially measured gene from a reference scRNA-seq data. Using these predicted values, RNA velocities can then be calculated for each cell in its spatial context, enabling the study of spatial differentiation trajectories.

To unlock the study of cellular differentiation dynamics in their spatial context, we propose SIRV (<u>S</u>patially Inferred <u>R</u>NA <u>V</u>elocity). SIRV is a pipeline representing a new use case of data integration of spatial and scRNA-seq data, combined with RNA velocity estimation for the spatial transcriptomics data. SIRV relies on domain adaptation to align spatial transcriptomics to matching scRNA-seq data. After alignment, the spliced and unspliced expressions of a spatially measured gene can be predicted from the neighboring scRNA-seq cells and used to calculate the corresponding spatial RNA velocity. In addition, SIRV can transfer various meta-data from scRNA-seq to spatial transcriptomics data, allowing, for example, detailed annotations of the spatial data. We used SIRV to study differentiation trajectories in the developing mouse brain and confirmed spatially localized trajectories at E10.5. Next, we applied SIRV to mouse organogenesis data and showed that SIRV can infer reproducible trajectories across different embryos. Finally, we verified our method by comparing the velocities inferred by SIRV to velocities inferred from measured spliced and unspliced counts in 10x Visium data of the developing chicken heart and MERFISH data from human osteosarcoma cells.

## 2. Methods

### 2.1. SIRV algorithm

The SIRV algorithm requires two inputs, the spatial transcriptomics data represented by a gene expression matrix, and the scRNA-seq data having three expression matrices corresponding to the spliced (mature mRNA), unspliced (immature mRNA) and full mRNA expression. The scRNA-seq data may also contain relevant metadata like cellular identity annotations, tissue/region of origin, etc. Using the set of shared genes between the two datasets, SIRV enriches the spatial transcriptomics data with spliced and unspliced expressions as predicted from the scRNA-seq. These spliced and unspliced expressions are then used to calculate the RNA velocity of each gene for each cell. Additionally, SIRV transfers the metadata from the scRNA-seq data to the spatial transcriptomics data. The SIRV algorithm consists of four major parts: (i) integration of the spatial transcriptomics and scRNA-seq datasets, (ii) predictions of spliced and unspliced expressions, (iii) label transfer (optional), and (iv) estimation of RNA velocities within the spatial context.

#### 2.1.1. Integration

The spatial transcriptomics and scRNA-seq datasets are integrated by finding the common signal between the two datasets. Similar to SpaGE^20^, the integration step is performed using PRECISE to define a common latent space^26^. In brief, using the set of shared genes across the two datasets, we calculate a separate Principal Component Analysis (PCA) for each dataset, and then align these separate principal components, resulting in principal vectors (PVs). These PVs have a one-to-one correspondence between the two datasets, and the highly correlated PV-pairs represent the common signal. Finally, both the spatial transcriptomics and scRNA-seq datasets are projected onto the PVs of the reference dataset (scRNA-seq in this case), producing an integrated and aligned version of both datasets.

This integration step is performed using the total (spliced + unspliced + ambiguous) mRNA expression matrix from the scRNA-seq side, together with the expression matrix of the spatial transcriptomics data. Thus, the spliced and unspliced expressions are only used in the (following) prediction step.

#### 2.1.2. Spliced and unspliced expression prediction

After obtaining the aligned datasets, SIRV enriches the spatially measured genes with spliced and unspliced expression predicted from the scRNA-seq data. Such prediction is performed using a kNN regression^20^. For each spatial cell *i*, we calculate the k-nearest-neighbors from the aligned scRNA-seq data and assign a weight to each neighbor inversely proportional to its distance.

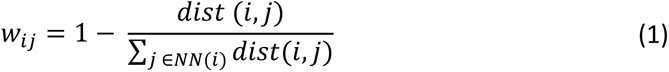

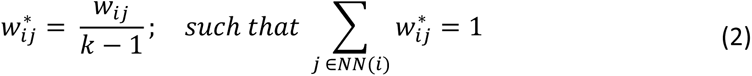

with *dist*(*i*,*j*) being the cosine distance between spatial cell *i* and scRNA-seq cell *j* ∈ *NN*(*i*), *k* equaling the number of nearest-neighbors used, and *w_ij_*^∗^ representing the weight between each spatial cell *i* and its *j*-th nearest neighbor.

For every spatially measured gene *g*, the spliced (*S_ig_*’) and unspliced (*U_ig_*’) expression are predicted by:

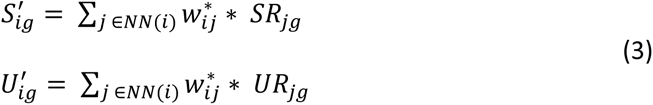

with *SR_jg_* and *UR_jg_* represent the spliced and unspliced expression of gene *g* from the scRNA-seq data, respectively.

#### 2.1.3. Label transfer

SIRV can annotate the spatial transcriptomics data with any relevant labels from the scRNA-seq data using the same kNN regression scheme as introduced earlier. Taking the cell identity annotation as an example: for each cell type *c* in the scRNA-seq annotation, we calculate a score *P*_*ic*_ expressing whether spatial cell *i* should be assigned to cell type *c* by aggregating the weights *w_ij_*^∗^ of the nearest neighbors annotated with *c*. The transferred cell type *C*_*i*_ for each spatial cell *i* is then selected based on the cell type with highest score:

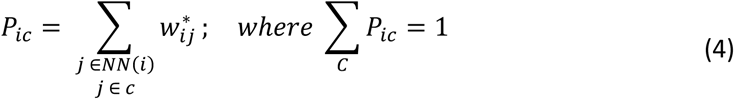

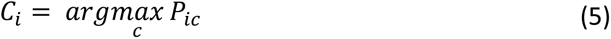

#### 2.1.4. RNA velocity analysis

After enriching the spatial genes with spliced and unspliced expressions, we applied the RNA velocity^5^ method (implemented in the scvelo^6^ python package) to investigate cellular development and differentiation. Following scvelo, first, we calculated the high-dimensional RNA velocity vectors for the spatially measured genes, next we projected and visualized these vectors on the spatial coordinates of the cells in order to define directions of cellular differentiation in the spatial context.

### 2.2. Datasets description

#### 2.2.1 Developing mouse brain

We used both spatial transcriptomics and scRNA-seq datasets from the Developing Mouse Brain Atlas^27^. Datasets were downloaded from http://mousebrain.org/downloads.html. The spatial transcriptomics data profiled the expression of 119 genes in an E10.5 mouse embryo, measured using the HybISS protocol^19^. Out of 25 different spatial slices provided by the authors, we selected the ’40 µm’ slice as it contains a clear structure of the brain. Cell segmentation was not provided with the data, however, we used the voxel version of the data (provided by the authors) which summarizes the spatial gene expression in a 2D grid of 30,000 pixels.

The scRNA-seq data profiled a developing mouse brain tissue from E7 to E18. To match the HybISS data, we only used E10 and E11 having a total 47,639 cells expressing 31,053 genes. Additionally, the scRNA-seq data was annotated with several metadata labels, we focused on the labels indicating the region (Forebrain, Midbrain and Hindbrain) and cellular identity. (‘Subclass’ annotation).

#### 2.2.2 Mouse organogenesis

We used three spatial datasets measured using the seqFISH protocol, representing three slices from the same mouse embryo^28^. The three datasets contained a total of 52,568 cells (19,451, 14,891 and 23,194 cells for Embryo1, Embryo2 and Embryo3, respectively) profiling the expression of 351 genes. Expression count data, spatial coordinates and corresponding metadata were downloaded from https://marionilab.cruk.cam.ac.uk/SpatialMouseAtlas/.

We chose the Gastrulation atlas^29^ as the reference scRNA-seq data for mouse organogenesis. We downloaded the complete atlas data from the MouseGastrulationData R package (https://bioconductor.org/packages/MouseGastrulationData/), then excluded cells with no celltype annotation, and selected only E8.5 to match the seqFISH spatial data with a total of 16,909 cells profiling 29,452 genes.

#### 2.2.3 Developing chicken heart

We obtained a pair of datasets from the developing chicken heart^8^, 10X Visium spatial data and 10X Chromium scRNA-seq data, from day 14 in the development. The provided count matrices by the authors did not contain the unspliced and spliced expression. For that we downloaded the fastq files from GEO (GSE149457), and used kallisto/BUStools^30,31^ to realign the reads to the reference genome (GRCg6a) while differentiating between intronic and exonic reads. Using the same list of cell/spot barcodes in the count matrices provided by the authors, we obtained 1,967 spots and 3,009 cells for the Visium and scRNA-seq data, respectively, both profiling 22,541 genes with the corresponding unspliced and spliced expression. Relevant metadata and spatial coordinates were downloaded from Github (https://github.com/madhavmantri/chicken_heart).

#### 2.2.4 Human osteosarcoma (U-2 OS)

We obtained three spatial transcriptomics datasets (batches) measured from human osteosarcoma (U-2 OS) cells using MERFISH^32^. Here, the spliced and unspliced expressions were replaced by cytoplasmic and nuclear expressions, respectively. We used batch 1 (645 cells) as our spatial data, while we concatenated batch 2 and 3 (400 and 323 cells, respectively) to act as simulated matching scRNA-seq data (ignoring the spatial locations of cells). We used the 9,050 genes detected with the non-overlapping probe design^32^.

### 2.3. Data preprocessing

First concerning the spatial datasets, the seqFISH and MERFISH datasets already had cellular resolution, the spots of the Visium data were treated as cells, while for the HybISS data, the pixels of the 2D grid were used as pseudo-cells. The gene expression of each pixel (pseudo-cell) is the count of the spots detected for each gene in that pixel location. To separate tissue from background, we filtered out any pseudo-cell with total counts across all genes less than 4. Next, we manually segmented only the brain tissue, ending with a total of 4,628 spatial pseudo-cells (Supplementary Fig. S1). Next, all spatial datasets were normalized by dividing the counts within each cell by the total count within that cell, multiplied by a scaling factor equal to the median number of counts across cells, and log(x+1) transformed.

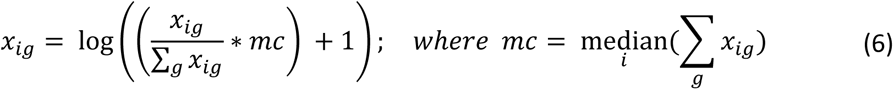

with *x_ig_* represents the expression of gene *g* in cell *i*, ∑*_g_ x_ig_* equals the total count in cell *i*, and *mc* is the median number of counts across cells. For the Visium data, we filtered out genes expressed in less than 10 cells, ending up with 12,295 genes, while no gene filtration was applied for the seqFISH and HybISS datasets.

Second concerning the scRNA-seq datasets, for the Gastrulation atlas, we renamed and regrouped few cell types based on previous recommendations^28^, and filtered out cell types with less than 25 cells. For the developing mouse brain data, cells having ‘Class’ annotation of ‘Bad cells’ or ‘Undefined’ were filtered out. Additionally, genes annotated as invalid (‘Valid’ = 0) were removed. Further, for all scRNA-seq datasets, we filtered out genes expressed in less than 10 cells. We ended up with 40,733 cells x 16,907 genes for the developing mouse brain data, 16,861 cells x 29,452 genes for the Gastrulation atlas, 3,009 cells × 10,143 genes for the 10X developing chicken heart data, while no genes were filtered out for the MERFISH simulated scRNA-seq data. Finally, each dataset was normalized by dividing the counts within each cell by the total count within that cell, multiplied by a scaling factor of 10^6^, and log(x+1) transformed.

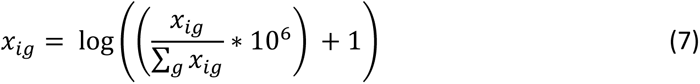

with *x_ig_* represents the expression of gene *g* in cell *i*, and ∑*_g_ x_*i*g_* equals the total count in cell *i*.

### 2.4. Weighted similarity metric

To quantitatively evaluate the estimated differentiation trajectories using SIRV, we compared the measured and estimated spatial/high-dimensional RNA velocity vectors using a weighted similarity metric score:

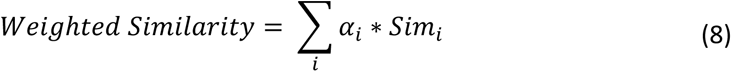

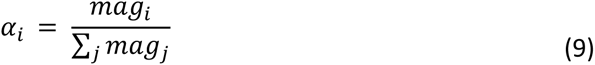

where *mag*_*i*_ is the magnitude of the spatial/high-dimensional velocity vector of spot *i*, *α*_*i*_ is the weight of spot *i* relative to its magnitude compared to all other spots, and *Sim*_*i*_ is the cosine similarity between the measured and estimated spatial/high-dimensional velocity vectors of spot *i*. Here, we emphasize the larger velocity vectors, which are more important to be estimated correctly compared to smaller velocity vectors.

### 2.5 RNA velocity gene contribution

For a selected set of cells *n*, we quantified the contribution of each gene in the resulting RNA velocity vectors, to observe which genes are most deriving the differentiation trajectory. First, we normalized the velocity vector of each cell to a unit vector. Next, the total contribution *C*_*g*_ of gene *g* is calculated as:

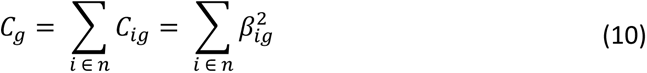

where *C*_*ig*_ is the contribution of gene *g* in cell *i*, which is equal to the square of the *g*-th component in the RNA velocity of cell *i* (*β*_*ig*_). The total contribution of each gene is obtained by summing the contributions along all cells in selection.

### 2.6. Implementation details

For the integration step we used 50 principal vectors^26^, and for the prediction step, we used *k* = 50^20^. For the RNA velocity analysis, we scaled the data to zero mean and unit variance features, next we calculate the top principal components (PCA) to build a neighborhood graph with 30 neighbors. Based on the percentage of explained variance in each dataset, we used 30, 50, 20 and 30 principal components for the HybISS, seqFISH, Visium and MERFISH datasets, respectively. This neighborhood graph is used to calculate a UMAP^33^ embedding of the data and cluster the data (in case of the HybISS and MERFISH data) using Leiden^34^ graph-based clustering (resolution = 1). Next, we calculate the RNA velocity vectors using the same number of principal components and neighbors. Finally, we project the high-dimensional velocity vectors on the UMAP coordinates and the spatial (x,y) coordinates using the *velocity_embedding* and *velocity_embedding_stream* functions. We used the scanpy^35^ python package (version 1.7.0) to perform data preprocessing, PCA, UMAP and Leiden clustering. While the scvelo^6^ python (version 0.2.3) package was used to calculate RNA velocities and their projections in the UMAP and the spatial context.

## 3. Results

### 3.1. SIRV overview

To enable RNA velocity estimation in spatial transcriptomic data, we developed SIRV (Fig. 1). Calculation of RNA velocity vectors requires measurements of mature (spliced) and immature (unspliced) mRNA expressions, which are missing in spatial transcriptomics data at the single-cell level. To overcome this limitation, we integrated the spatial transcriptomics data with matching scRNA-seq data, which includes spliced and unspliced mRNA expressions, from the same tissue (Methods). We have previously introduced SpaGE^20^, a method that integrates spatial and scRNA-seq datasets to predict whole-transcriptome expressions in the spatial data. Similar to SpaGE, SIRV relies on PRECISE^26^, a domain adaptation method, to correct for technical differences between the spatial and single-cell transcriptomic data. However, after integration, SIRV uses kNN regression to transfer (i.e. predict) the spliced and unspliced expressions for the spatially measured genes from the scRNA-seq data (Methods). The predicted spliced and unspliced expressions are used to calculate the RNA velocity vectors for every spatial cell. We project these vectors on the spatial coordinates of the tissue and derive flow fields by averaging the dynamics of spatially neighboring cells.

**Fig. 1.**
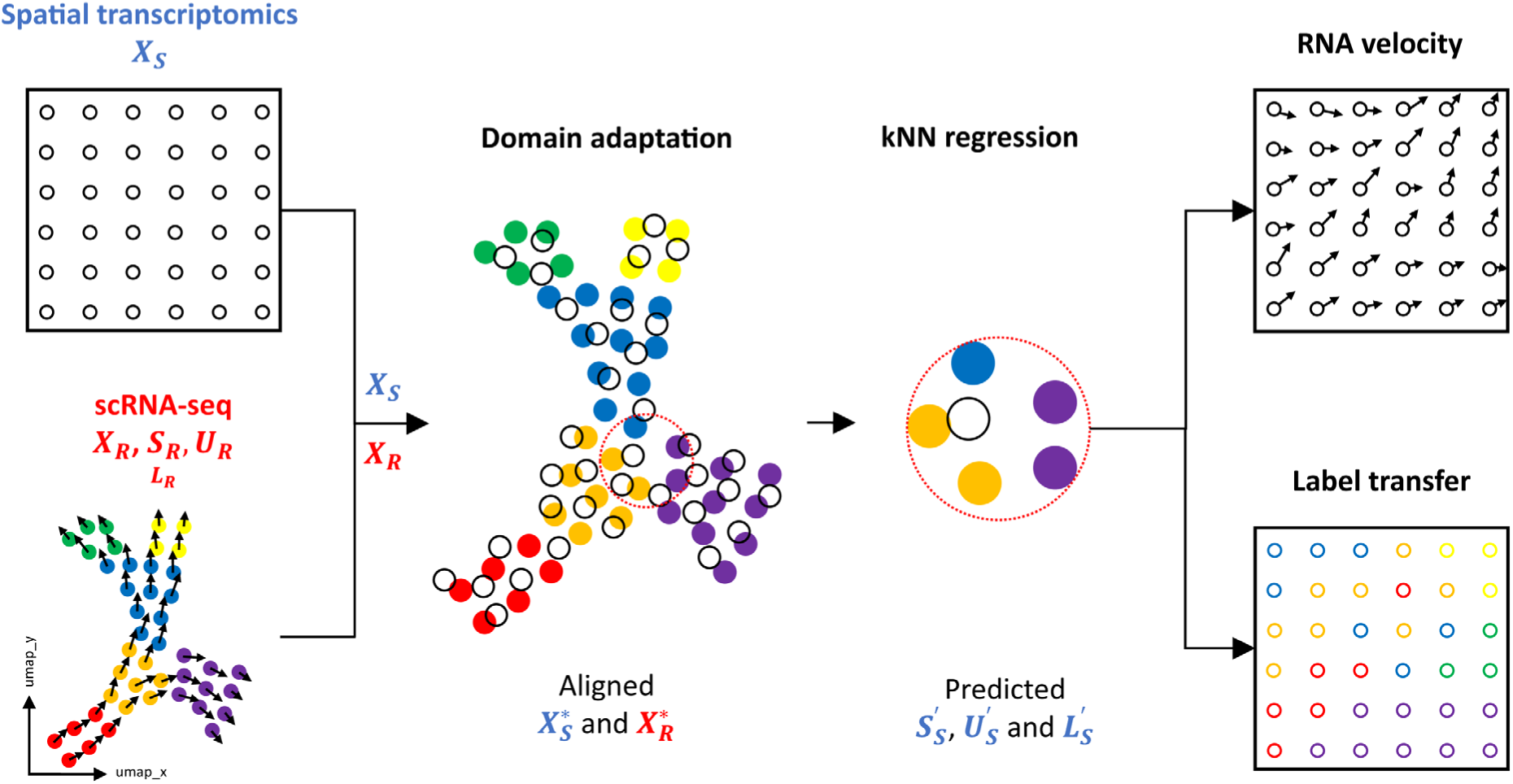
SIRV pipeline. SIRV takes as input a spatial transcriptomics data *X*_*S*_ and a reference scRNA-seq data. The latter contains spliced *S*_*R*_, unspliced *U*_*R*_ and total mRNA X_*R*_ expressions for each gene, and possibly label annotations *L*_*R*_ e.g. cellular identity. SIRV integrates the two datasets X_*S*_ and X_*R*_ using domain adaptation producing aligned datasets *X_S_*^∗^ and *X_R_*^∗^. Next, SIRV predicts the spatial spliced *S_S_*^′^ and unspliced *U_S_*^′^ expressions from the scRNA-seq data using kNN regression applied on *X_S_*^∗^ and *X_R_*^∗^. The predicted *S_S_*^′^ and *U_S_*^′^ expressions are used to calculate RNA velocity vectors, which are projected on the spatial coordinates of the tissue estimating spatial cellular differentiation trajectories. Additionally, SIRV transfers label annotations from scRNA-seq to spatial data (*L*_s_^′^) using the same kNN regression methodology.

Additionally, SIRV transfers various meta-data from the scRNA-seq to the spatial transcriptomics data using a similar kNN regression scheme (Methods). Considering cell identity labels, this label transfer feature offers an automated manner to annotate the spatial data. Since scRNA-seq captures the whole transcriptome, compared to a limited number of genes in the spatial data, the transferred annotations can represent more fine-grained cellular identities.

### 3.2. Reconstructing spatial differentiation trajectories in the developing mouse brain

To illustrate the utility of SIRV, we constructed spatial differentiation trajectories in the developing mouse brain. For this, we used spatial transcriptomic data mapping the expression of 119 genes in the E10.5 mouse brain using HybISS^27^. First, we clustered cells from the HybISS data into 21 clusters (Fig. 2A, Supplementary Fig. S2A). We mapped the cell clusters back to their spatial locations and observed that the clustering agrees with the spatial organization, i.e. the majority of the cell clusters are spatially-localized (Fig. 2B and Supplementary Fig. S2B-C), except for clusters 0, 15, 19 and 20. Furthermore, cluster 1 was spatially divided into two groups of cells, one localized to the midbrain and the other in the hindbrain.

**Fig. 2.**
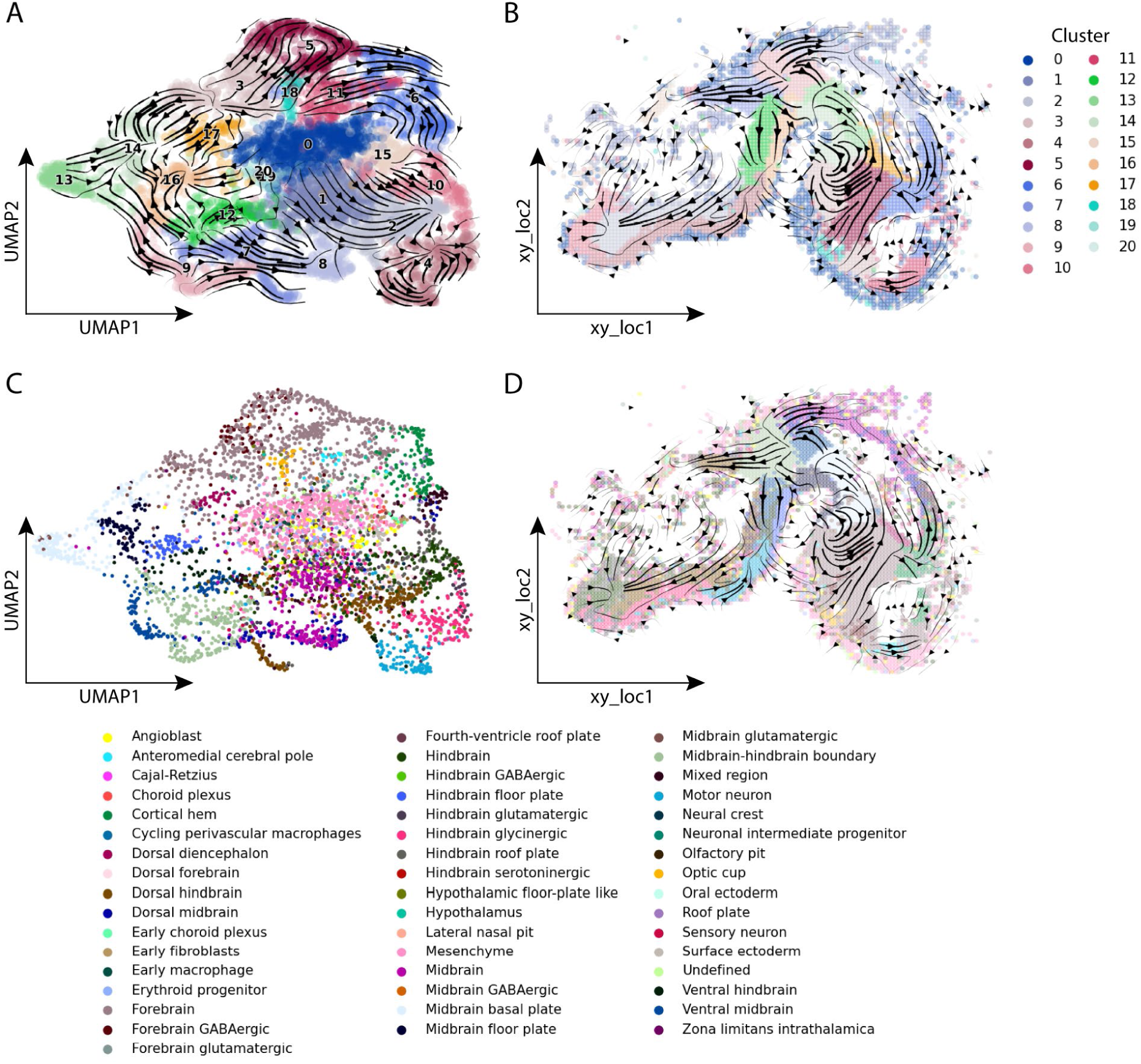
SIRV differentiation trajectories and label transfer for developing mouse brain data. **(A)** Main flow of SIRV predicted RNA velocities visualized by velocity streamlines, projected on the UMAP embedding of the HybISS spatial data, colored according to 21 cell clusters obtained using Leiden clustering. **(B)** Main flow of SIRV predicted RNA velocities visualized by velocity streamlines, projected on the spatial coordinated of the HybISS data, cell clusters show spatial localization. **(C)** UMAP embedding of the HybISS spatial data colored according to the ‘Subclass’ annotation transferred from the scRNA-seq data. **(D)** Spatial map showing the location of each ‘Subclass’ label in the tissue, with main flow streamlines of spatial RNA velocities as in **B**.

We integrated the HybISS spatial data with scRNA-seq data of the E10 and E11 mouse brain using SIRV and predicted the spliced and unspliced expressions for the spatially measured genes in every spatial cell. The two datasets shared 117 genes which were used for integration and prediction. With the predicted spliced and unspliced expressions for each of the 117 genes, we generated RNA velocity vectors for each spatial cell. The RNA velocity analysis showed differentiation patterns which are consistent with the HybISS clusters (Fig. 2A). For instance, cluster 9 differentiates into clusters 7, 8 and 13. Cluster 3 differentiates into clusters 5, 17 and part of 14, while the other part of cluster 14 is developed from cluster 13. Cluster 12 differentiates into cluster 16, while clusters 2 and 10 differentiate towards a common end point. Additionally, we observed that clusters 0, 15, 19 and 20 have close to zero magnitude velocity vectors. A possible explanation is that these clusters form the boundary of the brain (Supplementary Fig. S2C) and are not further involved in cellular differentiation at this stage of development.

Next, we projected and visualized the velocity vectors constructed by SIRV onto the spatial coordinates (Fig. 2B), which revealed the spatial differentiation dynamics of the cells. For example, cluster 9 is roughly located at the midbrain-hindbrain boundary, and the velocity vectors branches in 3 directions towards clusters 7, 8 and 13. To obtain a more detailed view of the spatial RNA velocities, we visualized the spatial velocity vectors at the single-cell level (Supplementary Fig. S3A). Results show that the velocity vectors follow consistent spatial paths across the different cell clusters and different brain regions. If we consider cluster 9 again, the spatial velocity vectors at the cell level show the same differentiation into 3 branches (Supplementary Fig. S3B). Furthermore, we can clearly observe the differentiation of cluster 12 into 16, and cluster 13 into a part of cluster 14 (Supplementary Fig. S3C). Moreover, cluster 2 forms a relatively long path of differentiation through the Hindbrain (Supplementary Fig. S3D), reaching a common end point together with cluster 10. Interestingly, investigating spatial differentiation trajectories revealed patterns which were not apparent in the UMAP (Fig. 2A). For instance, the spatial velocity vectors suggest that cluster 3, located in the forebrain, differentiates towards cluster 5 and part of 14. However, cluster 3 indirectly differentiates to cluster 17 through cluster 5 (Supplementary Fig. S3E). Moreover, from the spatial context, we observed that cluster 5 further differentiates to cluster 6. Together, these results show the potential of SIRV to identify spatial differentiation trajectories that cannot be obtained from dissociated data representation alone.

### 3.3. Interpretation of spatial RNA velocities in the developing mouse brain

To gain additional insights into the spatial trajectories, we used SIRV to transfer cell annotations from the scRNA-seq to the HybISS spatial data. First, we transferred the brain region annotation (forebrain, midbrain and hindbrain) from the scRNA-seq to the HybISS data. In the UMAP embedding of the HybISS spatial data, the three brain regions form three different groups with some overlap in the middle (Supplementary Fig. S4A). This overlap can be explained by the group of mesenchyme cells forming the border of the brain (Fig. 2C-D). When visualized in their spatial coordinates, we observe that the regional annotation of cells is in agreement with the anatomical structure of the brain (Supplementary Fig. S4B and S1C-D). Next, we transferred the ‘Subclass’ annotations which formed distinct groups in the UMAP embedding of the spatial data (Fig. 2C). Spatially, the obtained Subclass labels also showed a well-separated and structured spatial organization (Fig. 2D and Supplementary Fig. S5). Subclasses belonging to either hindbrain, midbrain or forebrain clusters are found in their respective brain regions with little overlap to neighboring regions, validating the estimation of these labels. For example, midbrain-hindbrain boundary cells are positioned around the isthmus, and the cortical hem and dorsal diencephalon cells localize in the forebrain as expected^27,36^. The scRNA-seq dataset has a total of 104 ‘Subclass’ label annotations. Using SIRV, only 49 labels were transferred to the spatial data. These labels ranged in size from 18 to 5065 cells in the scRNA-seq dataset, while the other 55 non-transferred labels ranged from 1 to 157 cells (Supplementary Fig. S4C). This suggests that SIRV is able to transfer the relevant annotations to the spatial data, including small cellular populations in the corresponding scRNA-seq dataset, and it’s not only biased towards transferring the large abundant cell populations.

We visualized the spatial RNA velocity vectors at the single-cell level together with the transferred ‘Subclass’ annotations (Fig. 3A). The midbrain-hindbrain boundary cells (cluster 9) show three branched trajectories, differentiating towards midbrain, ventral midbrain and dorsal hindbrain cells (Fig. 3B). Comparing the 21 cell clusters with their ‘Subclass’ annotation indeed shows that cluster 9 mostly maps to the midbrain-hindbrain boundary subclass (Supplementary Fig. S4E). In the midbrain, midbrain basal plate cells differentiate into midbrain floor plate cells (Fig. 3C). In the hindbrain, dorsal hindbrain cells differentiate towards hindbrain cells (Fig. 3D). Considering the forebrain region, cells annotated as forebrain (mostly covering clusters 3, 5 and 11) differentiate into dorsal diencephalon (cluster 17) and cortical hem (cluster 6) (Fig. 3E and Supplementary Fig. S4E). While cluster 0, 15, 19 and 20 are mostly composed of mesenchyme cells forming the borders of the brain (Supplementary Fig. S4E), no differentiation was associated with these clusters in agreement with mesenchyme cells which also have almost zero magnitude RNA velocity vectors. To evaluate the trustworthiness of the generated velocities, we calculated the velocity confidence scores from the scvelo package. We observed relatively high confidence scores across the majority of the brain regions (Supplementary Fig. S4D), with an average confidence score of 0.87 across all cells.

**Fig. 3.**
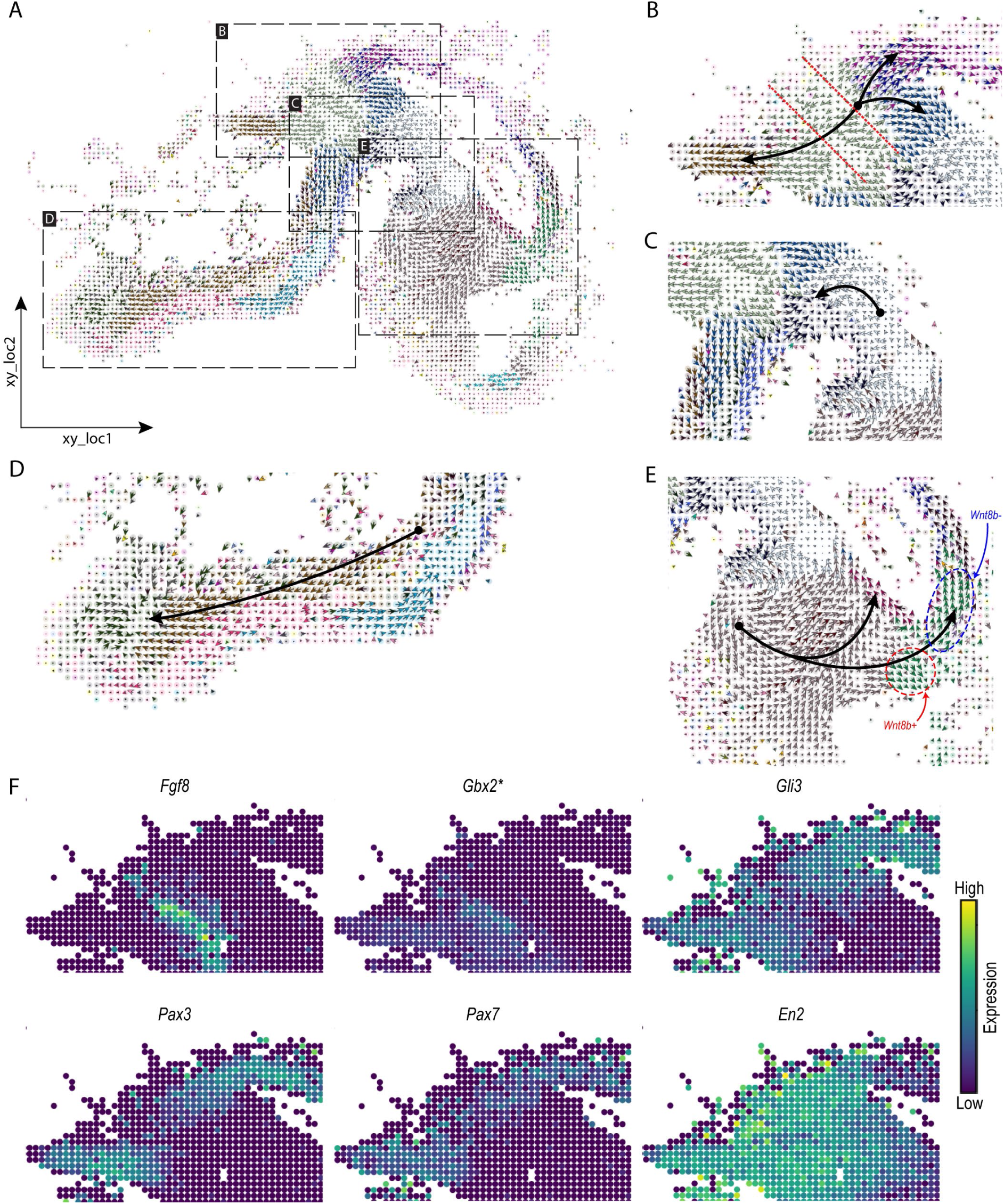
**(A)** Cell-level RNA velocities projected on the spatial coordinates of the HybISS spatial data, colored according to the ‘Subclass’ annotation transferred from the scRNA-seq data (same legend as Fig. 2C-D). **(B-E)** Zoom-in views on the same spatial differentiation trajectories as in Fig. 3; **(B)** midbrain-hindbrain boundary, dashed-lines highlight the isthmus region, **(C)** part of midbrain, **(D)** hindbrain, and **(E)** forebrain, highlighting two cortical hem populations (Wnt8b+ in red, and Wnt8b-in blue). Black arrows show branching or linear spatial differentiation directions between subclasses (these arrows are drawn manually to highlight certain differentiation trajectories). **(F)** Overlaid expression of marker transcription factors following the differentiation trajectory in the hindbrain and midbrain. *Predicted expression from scRNA-seq data using SpaGE^20^.

Next, we focused on the spatial differentiation trajectory at the midbrain-hindbrain boundary and aimed to confirm the trajectory suggested by these spatial RNA velocity vectors (Fig. 3B). The isthmus is an important organizer during midbrain and hindbrain development and is marked by the expression of *Fgf8*^37^. *Otx2* expression rostral to the isthmus and *Gbx2* expression caudal to the isthmus restrict *Fgf8* expression. In addition, *Gli3* is implicated as a regulator of *Fgf8* expression and is essential for establishing the tectum, isthmus and cerebellum during early development^38^. The combinatorial expression of *Otx2*, *Pax2/5* and *En1/2* drives cells towards a mesencephalic fate, and *Pax3/7* expression further drives differentiation towards the tectum^37,39^. *Fgf8, Pax3/7,* and *En2* expression was detected in their expected regions in the isthmus and midbrain (Fig 3F). Examining the velocity plot of *Gli3* indeed shows the midbrain-hindbrain boundary cells as an earlier differentiation stage, while dorsal hindbrain cells and midbrain/dorsal midbrain cells (tectum) are late stages (Supplementary Fig S6A). Similar observations can be obtained from *Pax3/7*, while *En2* shows the ventral midbrain cells as a later differentiation stage following the midbrain-hindbrain boundary cells. We repeated the RNA velocity analysis using only the five genes (*Fgf8*, *Gli3*, *Pax3*, *Pax7* and *En2*) involved in the midbrain-hindbrain differentiation trajectory. We were able to partially reconstruct the spatial trajectory obtained using all spatial genes (Supplementary Fig. S6B) with a weighted cosine similarity of 0.33 (Methods). Although the direction of the ventral midbrain cells is changed, the branching of the midbrain-hindbrain boundary cells into dorsal hindbrain and midbrain/dorsal midbrain cells could still be observed.

We then asked whether *Pax3/7* were also the main contributors to the RNA velocity trajectory of the cells differentiating into the tectum (Methods). For the cells differentiating towards the tectum, the top contributing genes were *Gli3, Egfem1,* and *Pax3,* in agreement with the previously described role of *Gli3* and *Pax3* in the development of the tectum. However, *Egfem1* (EGF-like and EMI domain-containing protein 1), has not previously been implicated in neural development of the tectum and its function within the central nervous system is still unknown^40^.

Furthermore, we noticed that SIRV predicted two distinct trajectories in the cortical hem (Fig. 3E). The cortical hem is important for the formation of the choroid plexus and hippocampus, and gives rise to Cajal-Retzius cells^41^. The development of the cortical hem is impaired in Gli3-deficient mice^42,43^. *Lmx1a* is required to repress *Lhx2* expression in the cortical hem, preventing progenitors from adopting a hippocampal fate and ensuring the production of Cajal-Retzius neurons^44^. The Wnt family member *Wnt8b* marks the cortical hem^45,46^ and *Wnt8b* expression appeared to divide the trajectories in a *Wnt8b^+^*trajectory rostrally and a *Wnt8b^-^* trajectory dorsally (Fig. 3E). We therefore asked whether these genes contribute to the RNA velocity trajectories in the cortical hem, and whether *Wnt8b* defines the rostral trajectory. In contrast to the RNA velocity trajectory in the dorsal midbrain, the gene contribution to the trajectories of the cortical hem were more dispersed across several genes. The top contributing genes for the two subsets showed considerable overlap (*Gli3, Lmx1a*) and clear differences (*Unc5c, Pax6,* in *Wnt8b^-^*, *Eya4 in Wnt8b^+^*). Surprisingly, *Wnt8b* barely contributed to the direction of the trajectory in either subset of cells. *Pax6* is involved in cortical development, but lowly expressed in the cortical hem^47^. *Unc5c (*Unc5 Netrin Receptor C), a receptor to the axon guidance cue netrin-1, and *Eya4* (EYA transcriptional coactivator and phosphatase 4) have not been linked to early cortical hem development before. Conclusively, SIRV shows reliable RNA velocity trajectories in the developing mouse brain and could implicate novel genes in the differentiation of cell types during neural development.

### 3.4 SIRV reveals spatial differentiation trajectories that are missed using scRNA-seq only

Next, we wondered if inferring spatial trajectories using SIRV reveals patterns that cannot be captured using RNA velocity from the scRNA-seq data alone. We applied RNA velocity to the scRNA-seq data of the developing mouse brain, using the top 2,000 highly-variable genes (HVGs). We could not detect the differentiation trajectory of the midbrain-hindbrain boundary cells (Fig. 4A), which is the clearest trajectory captured by SIRV (Fig. 3B). We then wondered if the results are driven by the highly selected genes measured in the HybISS data. To test the effect of gene selection on velocity estimation, we inferred velocities from the scRNA-seq data using the same 117 genes measured in the HybISS dataset. Yet, the midbrain-hindbrain boundary trajectory was not captured (Fig. 4B). To make sure that such differentiation trajectory was not over-taken by a stronger signal in the scRNA-seq data, we selected the cell populations involved in the midbrain-hindbrain boundary differentiation trajectory and applied the RNA velocity to these cells only. Again, the trajectory was not captured, neither using the 2,000 HVGs nor the 117 spatial genes (Supplementary Fig. S7A-B).

**Fig. 4.**
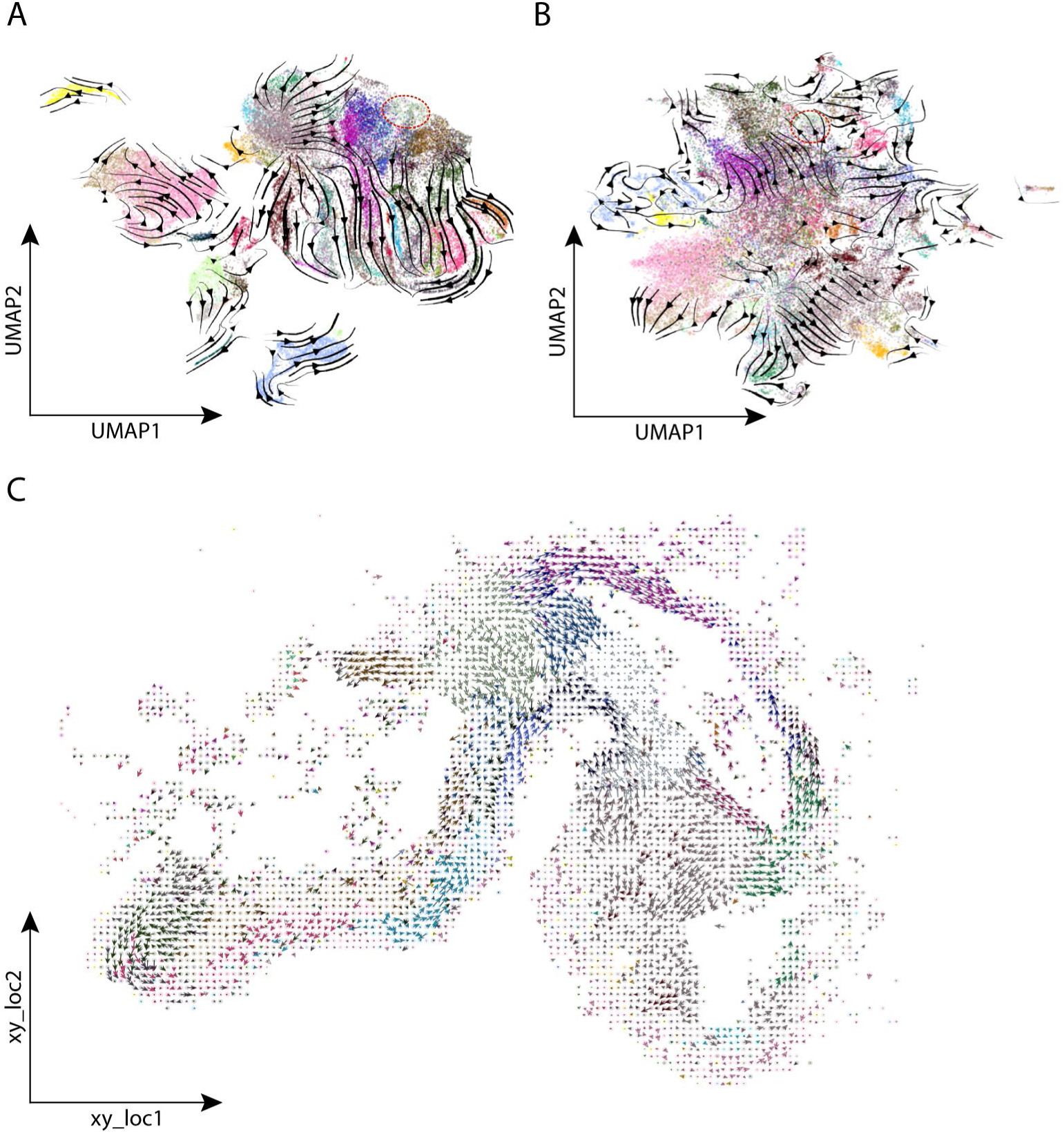
**(A-B)** Main flow of RNA velocities visualized by velocity streamlines projected on the UMAP embedding of the scRNA-seq data of the developing mouse brain, RNA velocity was calculated using **(A)** top 2,000 HVGs, or **(B)** 117 spatial genes. Cells are colored according to the ‘Subclass’ annotation of the scRNA-seq (same legend as Fig. 2C-D). The branching trajectory of the midbrain-hindbrain boundary cells (highlighted in red) is not detected in both cases. **(C)** Cell-level RNA velocities projected on the spatial coordinates of the HybISS spatial data, where SIRV was used to predicted transcriptome-wide spliced and unspliced expressions, and RNA velocity analysis was performed using the top 2,000 HVGs. Cells are colored according to the ‘Subclass’ annotation transferred from the scRNA-seq data (same legend as Fig. 2C-D).

### 3.5 Relying on spatially measured transcriptomes provides better spatial differentiation trajectories

Although the spliced and unspliced counts of individual genes are predicted by SIRV for every spatial location, we do use the measured gene count when calculating the (spatial) RNA velocities. We decided to do so because the spatial transcriptomics data has a higher detection rate (fraction of cells expressing a certain gene) (Supplementary Fig. S7C). To test the validity of this choice, we compared the measured spatial transcriptomics data (X_*s*_) with the sum of the predicted spliced (*S*_s_^′^) and unspliced (*U*_s_^′^) expressions as derived from the scRNA-seq data. We indeed observed high correlations, comparable to the correlation between scRNA-seq data (X_*r*_) and the measured spliced (*S*_*r*_) and unspliced (*U*_*r*_) expressions (Supplementary Fig. S7D). To further test that the use of the spatially measured gene counts is to be preferred over predicted ones, we generated spatial RNA velocities using the summed predicted spliced and unspliced expressions as gene counts. Supplementary Fig. S7E indeed shows that then the predicted spatial trajectories do deteriorate when compared to using the spatially measured gene counts.

SIRV is capable of predicting the spliced and unspliced expressions transcriptome-wide, not only for the spatially measured genes. We thus were interested whether calculating the spatial RNA velocities based on a full (predicted) transcriptome would outperform one in which the RNA velocities were only based on the measured spatial genes. Hereto, we predicted with SIRV the spliced and unspliced expressions for all genes from the scRNA-seq. The spatial gene counts are now represented by the summation of the predicted spliced and unspliced expressions (X′_*s*_ = *S*^′^ + *U*^′^) as we don’t have (spatial) measurements for most genes. Using the top 2,000 HVGs, we performed the RNA velocity and projected the velocity vectors on the spatial coordinates of the mouse brain, obtaining consistent spatial differentiation paths across different brain regions (Fig. 4C). However, the spatial differentiation trajectories obtained using the 117 spatial genes with their corresponding measured spatial data (Fig. 3A) were more clearly defined, showing the value of using the spatial data even with smaller set of genes.

### 3.6 Reproducible spatial differentiation trajectories in mouse organogenesis

To illustrate the versatility of SIRV, we next analyzed spatial transcriptomic data of mouse organogenesis^28^. This data mapped the spatial expression of 351 genes from three mouse embryos at embryonic day (E)8.5–8.75 using seqFISH. As a reference scRNA-seq, we used the Gastrulation scRNA-seq atlas (E8.5)^29^. Based on 348 shared genes, we integrated the two datasets and predicted the spliced and unspliced expressions of those shared genes for each spatial cell. Next, for each of the three spatial embryos, we calculated the RNA velocity vector of each cell, then projected and visualized these vectors onto the spatial coordinates. For embryo 1, SIRV estimated several spatial differentiation trajectories across different cell types (Fig. 5A). For example, the gut tube cells differentiate anterio-dorsally (Fig. 5B), while the mixed mesenchymal mesoderm cells are differentiating towards splanchnic mesoderm cells (Fig. 5C). These examples, in addition to the full differentiation spectrum, were reproducibly estimated by SIRV across the three embryos at the single-cell level (Fig. 5B-C, Supplementary Fig. S8). In terms of trustworthiness of these spatial velocities, we obtained high average velocity confidence scores of 0.92, 0.90 and 0.89 across the three embryos, respectively (Supplementary Fig. S9).

**Fig. 5.**
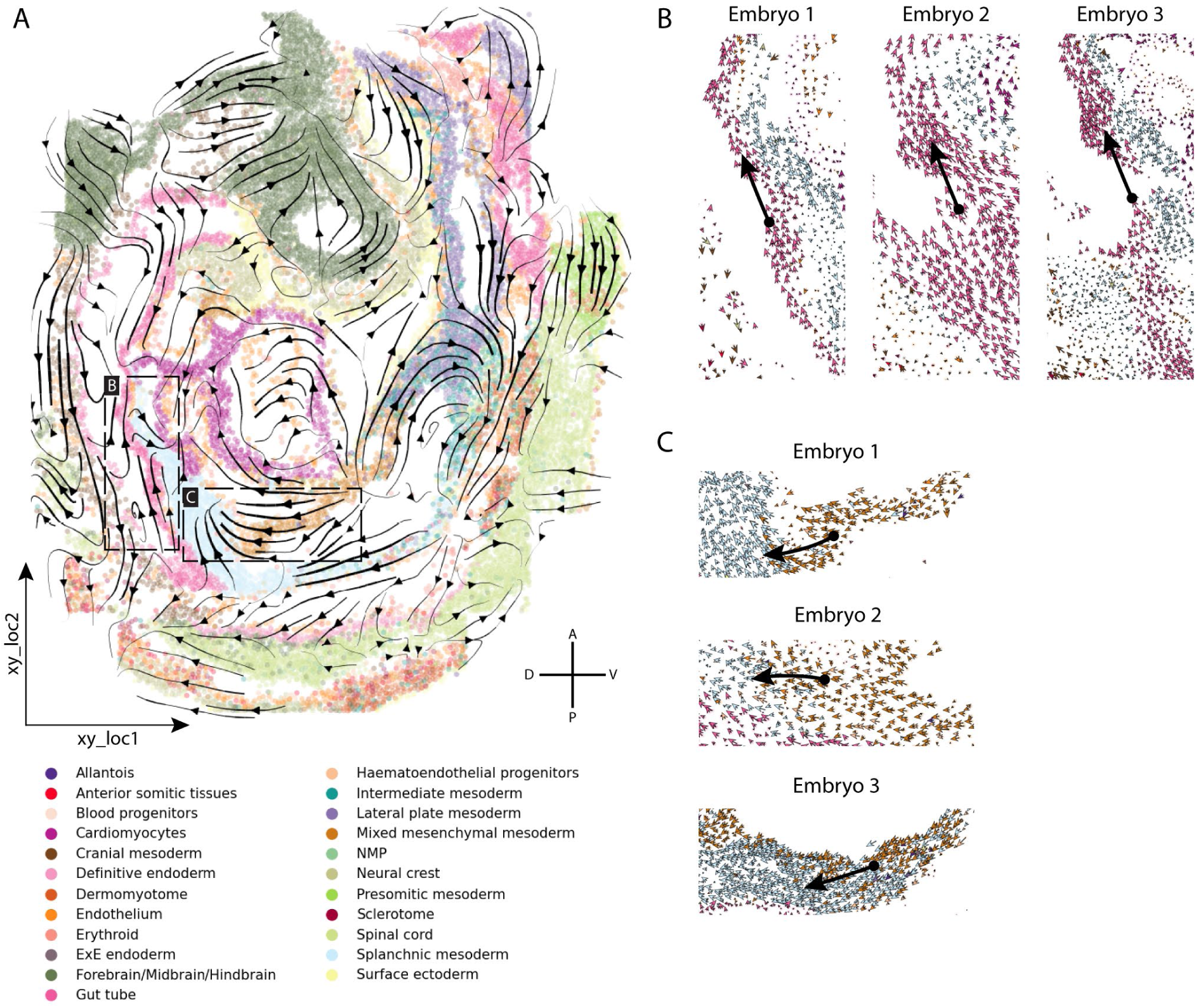
SIRV spatial differentiation trajectories for mouse organogenesis. **(A)** Main flow of the SIRV predicted RNA velocities across different cell types, visualized by velocity streamlines, projected on the spatial coordinates of *Embryo 1* from the mouse organogenesis SeqFISH data. **(B-C)** Zoom-in views on the cell-level spatial RNA velocities, showing reproducible results across the three embryos, where in **(B)** gut tube cells differentiate in the Anterior-Dorsal direction, while in **(C)** mixed mesenchymal mesoderm cells differentiate towards splanchnic mesoderm cells (Black arrows are drawn manually to highlight certain differentiation trajectories).

Furthermore, we used SIRV to transfer the cell type annotations from the scRNA-seq to the seqFISH data. Since the seqFISH data was previously annotated in a similar manner, we used the seqFISH data annotation as ground truth to evaluate the accuracy of the label transfer. Our results show accurate annotation of various cell types including cardiomyocytes, erythroid and brain cells, among others (Supplementary Fig. S10A), while other cell types, such as blood progenitors and ExE endoderm, were more challenging to annotate. SIRV obtained an overall accuracy of 0.74 and a median F1-score of 0.69 across all 23 cell types. Of note, annotations of the seqFISH data were refined manually by the authors, which partially explains the misclassification of cells by SIRV^28^. To test this observation, we compared the performance of label transfer between SIRV, Seurat (CCA+MNN)^22^ and Tangram^48^, where the latter is reported as the top performing integration method for spatial and scRNA-seq data^25^. Seurat obtained a comparable annotation performance with an accuracy of 0.70 and a median F1-score of 0.61 (Supplementary Fig. S10B), while Tangram showed lower performance in comparison to SIRV and Seurat, with an accuracy of 0.53 and a median F1-score of 0.41 (Supplementary Fig. S10C).

Next, we evaluated the accuracy of the integration step of SIRV, for that we benchmarked against Harmony^49^ and Seurat^22^, top performing methods for integration of single-cell transcriptomics data^50^. The UMAP^33^ embeddings of the integrated seqFISH and scRNA-seq data are comparable between SIRV, Harmony and Seurat (Supplementary Fig. S11). For a quantitative comparison, we calculated the average silhouette score (ASW) and the Local Inverse Simpson’s Index (LISI)^49^ using both batch and cell type identifiers across all methods. SIRV integration was comparable to Harmony and Seurat. For cell type separation, SIRV obtained the overall best cell type ASW (higher is better) of 0.145 compared to 0.119 and 0.101 for Harmony and Seurat, respectively. However, SIRV ranked 2nd in terms of cell type LISI (optimal is 1) of 1.42 compared to 1.44 and 1.39 for Harmony and Seurat, respectively. For batch mixing, SIRV obtained the overall best batch ASW (lower is better) of 0.001 compared to 0.002 and 0.006 for Harmony and Seurat, respectively. However, SIRV ranked last in terms of batch LISI (optimal is 2) of 1.28 compared to 1.38 and 1.37 for Harmony and Seurat, respectively.

Taken together, these results illustrate that SIRV can estimate robust spatial differentiation trajectories, and accurately integrate and annotate spatial data using reference scRNA-seq data.

### 3.7. Verification of SIRV using Visium and MERFISH data

To verify the spatial RNA velocity vectors estimated using SIRV, we used a 10x Visium data of the developing chicken heart^8^. 10x Visium is a sequencing-based method, allowing the quantification of spliced and unspliced mRNA reads in 55 µm wide tissue spots composed of multiple cells. We used kallisto and bustools^30,31^ to quantify the spliced and unspliced counts from the 10x Visium data as well as from a matching scRNA-seq data from the same study^8^. We used the measured 10x Visium spliced and unspliced expressions to calculate RNA velocity vectors (hereafter: measured velocities), showing various spatial differentiation trajectories when projected to the spatial coordinates of the tissue (Fig. 6A). Next, we used SIRV to integrate the 10x Visium data with the 10x scRNA-seq data and predicted the spliced and unspliced expressions for each spatial spot from the scRNA-seq data. We used these predicted spliced and unspliced expressions to calculate the RNA velocity vector of each spot (hereafter: estimated velocities), which we then also projected on the spatial coordinates (Fig. 6B). We observed a general agreement between the measured and estimated velocity vectors at the single-spot level across different regions of the tissue (Fig. 6A-B, Supplementary Fig. S12A-B), including the erythrocytes, fibroblasts and cardiomyocytes in the left ventricle. To quantitatively evaluate the estimated spatial velocities, we calculated the cosine similarity between the measured and estimated spatial velocity vectors (Methods, Supplementary Fig. S12C-D). We obtained relatively high similarity at different regions of the tissue including the examples previously mentioned, with an overall weighted similarity of 0.45 between the high-dimensional velocity vectors and 0.25 between the 2D spatial velocity vectors across the whole tissue. It is important to note that the measured and estimated RNA velocity vectors cannot be directly compared, since the measured velocities are calculated from multiple cells captured by each spot in the 10x Visium data, while the estimated velocities are calculated based on individual cells from the scRNA-seq data.

**Fig. 6.**
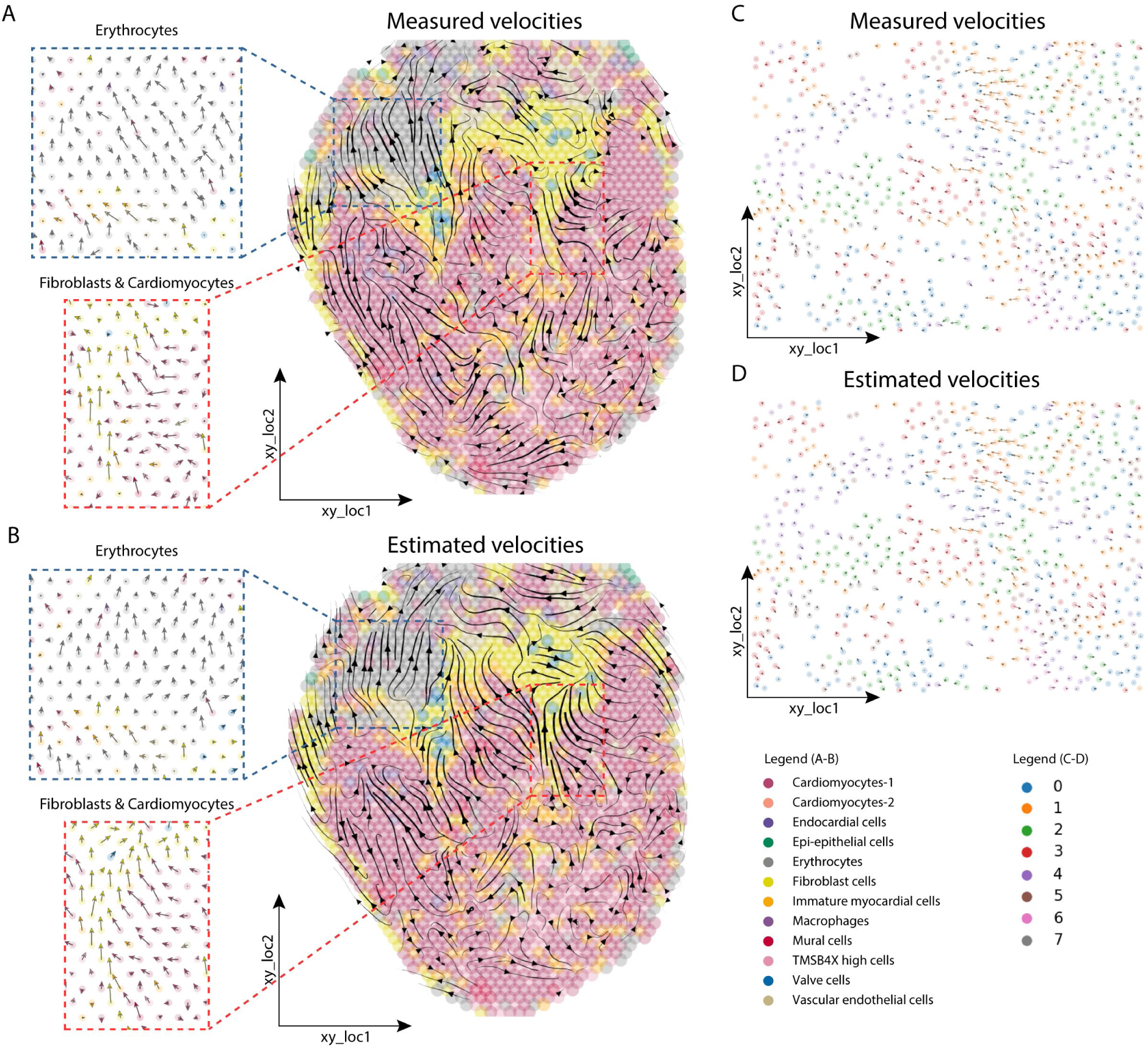
Validation of estimated spatial differentiation trajectories using SIRV. **(A-B)** Main flow of the **(A)** measured and **(B)** SIRV estimated spatial RNA velocities across different cell types, visualized by velocity streamlines, projected on the spatial coordinates of the developing chicken heart 10X Visium dataset. Various regions show high agreement between measured and estimated spatial differentiation trajectories, including the erythrocytes (blue box) and fibroblasts and cardiomyocytes in the left ventricle (red box). Zoom-in views show similar agreement at the single-spot level. **(C-D)** Cell-level RNA velocities of the **(C)** measured and **(D)** SIRV estimated velocities projected on the spatial coordinates of the human osteosarcoma MERFISH dataset, showing high agreement. Cells are colored according to clusters obtained using Leiden clustering.

To overcome the limited spatial resolution of the 10x Visium data, we turned to MERFISH data from human osteosarcoma (U-2 OS) cells^32^. MERFISH is a subcellular resolution imaging-based method that detects the spatial location of mRNA molecules in a cell. While it’s not possible to distinguish spliced and unspliced mRNAs using MERFISH, cytoplasmic and nuclear mRNA can be used instead^32^. Here the assumption is that spliced reads are more enriched in the cytoplasm while the unspliced reads are more enriched in the nucleus. First, we clustered the MERFISH data into 8 clusters, then we used the measured cytoplasmic (spliced) and nuclear (unspliced) expressions to calculate RNA velocity vectors (i.e. the measured velocities). We then simulated matching scRNA-seq data by ignoring the spatial location of cells from other batches of the MERFISH data (Methods). Using SIRV, we then set out to predict the cytoplasmic and nuclear expressions for each spatial cell from the simulated scRNA-seq data (i.e. the estimated velocities). Projecting the measured and estimated velocities on the spatial coordinates of the cells showed a high agreement at the single-cell level (Fig. 6C-D). Since the MERFISH data contains only one cell type, the identified cell clusters represent different states in the cell-cycle, which are captured by the measured velocities as was previously shown^32^ (Supplementary Fig. S13A). The estimated velocities using SIRV similarly reconstruct the cell-cycle process captured by the measured velocities (Supplementary Fig. S13B). Overall, we obtained a weighted similarity of 0.99 between the high-dimensional velocity vectors and 0.84 between the 2D spatial velocity vectors (Supplementary Fig. S13C-D).

Further, we benchmarked the prediction of spliced and unspliced expressions using SIRV against Harmony and Seurat. While Seurat can perform the whole pipeline of integration and prediction of new genes, Harmony only provides the integration of spatial and scRNA-seq data but lacks a method to predict the expression of new genes. Therefore, we applied the same kNN regression scheme as used in SIRV to predict spliced and unspliced expressions. We calculated the Spearman correlation between the original spatially measured and predicted spliced and unspliced expressions across all methods. For the 10x Visium data, all methods performed similarly with a median Spearman correlation of 0.03, 0.02 and 0 for SIRV, Harmony and Seurat, respectively, for the spliced data (Supplementary Fig. S14A). While all methods had a median correlation of 0 for the unspliced data (Supplementary Fig. S14B), showing the challenge of matching spot data (being a collection of cells) with single-cell data. For the MERFISH data, SIRV showed improved performance with a median correlation of 0.21 for both spliced and unspliced data, compared to 0.08 and 0 for Harmony and Seurat, respectively (Supplementary Fig. S14

C-D). Additionally, similar to SIRV, we used the predicted spliced and unspliced expressions from Harmony and Seurat and performed the RNA velocity analysis. Comparing the measured and estimated RNA velocities shows that SIRV outperforms both Harmony and Seurat on both 10x Visium and MERFISH datasets. For the 10x Visium, Harmony and Seurat respectively obtained a weighted similarity of 0.33 and 0.40 between the high-dimensional velocity vectors, and 0.04 and 0.16 between the 2D spatial velocity vectors. While for the MEFRISH data, Harmony and Seurat respectively obtained a weighted similarity of 0.82 and 0.01 between the high-dimensional velocity vectors, and 0.56 and 0.08 between the 2D spatial velocity vectors.

Taken together, these verification results show that SIRV-estimated spatial differentiation trajectories resemble those that could be obtained directly from measured spatial data at the single-cell or spot level.

## 4. Discussion

We developed a computational method to transfer RNA velocities from scRNA-seq data to spatial transcriptomics data at the single-cell level, allowing the investigation of differentiation trajectories in their spatial context. Our method, SIRV, can be used to enrich any spatial transcriptomics data with spliced and unspliced mRNA expressions, as well as any additional cell meta-data, from a matching scRNA-seq data. We have shown that exploiting the spatial data is valuable and reveals differentiation patterns that are not captured by the scRNA-seq data only.

Since the HybISS spatial data was not originally annotated with cellular identities, the spatial differentiation trajectories obtained with SIRV could not be easily interpreted using the spatial data as is, but this was overcome by transferring cell annotations with SIRV. However, the interpretation of the estimated spatial velocity vectors is still challenging, i.e. the differentiation directions might only imply that cells differentiate from one state to another, or also imply that cells do migrate in the tissue during development where RNA gradient might be necessary^51,52^.

SIRV estimated reproducible spatial differentiation trajectories in mouse organogenesis data from three different embryos. Additionally, SIRV-estimated RNA velocities showed consistency with RNA velocities calculated from the measured spatial data. When applied on the 10x Visium developing chicken heart data, the spatial velocities estimated using SIRV resembled the measured velocities at the single-spot level. However, this comparison suffers from the fact that the 10x Visium measures mRNA content in 55 µm wide spots, composed of a few cells, while the corresponding scRNA-seq captures single cells, which could explain the overall lower similarity. In contrast, applying SIRV on the MERFISH data, in which we simulated scRNA-seq data from different MERFISH batches, yielded a high similarity, being a proof-of-concept that SIRV is able to correctly reproduce the measured differentiation trajectories. In addition, we observed regions with poorly estimated spatial velocities in the 10x Visium data, having even negative cosine similarities (Supplementary Fig. S12D). These negative similarities are still obtained for the MERFISH data (Supplementary Fig. S13D), which shows that SIRV is not always able to estimate the RNA velocity accurately. However, these negative similarities are much less present for the MERFISH data in comparison with the 10x Visium data, which can be explained by matching spots with single-cells instead of cells with cells as done in this MERFISH experiment.

In the current study, the spatial information is not directly used during the RNA velocity calculation and is mainly used to project and visualize the RNA velocity vectors in the spatial domain. However, the spatial information is implicitly driving the definition of the RNA velocities between cells, as the velocity graph calculation is based on the measured spatial gene expression, which tends to be more similar for closeby cells. Further development could benefit from explicitly using the spatial information to inform RNA velocity calculation.

With SIRV, we generalized the idea of transferring any relevant features from scRNA-seq to spatial data. This can be further extended to other applications, for instance, transferring the expression of different RNA species measured using VASA-seq^53^. Furthermore, in this study we used the steady-state model from scvelo^6^ to calculate the RNA velocity. SIRV can also be extended to other upcoming models, that require spliced and unspliced expressions, developed to overcome current limitations such as the lack of modeling of transcriptional bursts^54^. However, different RNA velocity methods need to be carefully evaluated to ensure a proper comparison on their spatial equivalence.

SIRV was able to estimate biologically relevant spatial differentiation trajectories in the developing mouse brain data. However, it is important to note that projecting the high-dimensional RNA velocity vectors into two-dimensional coordinates may introduce undesired artifacts, as cells are forced to point towards nearby cells. Therefore, novel estimated differentiation trajectories require proper validation.

Concluding, SIRV produces valuable spatial differentiation trajectories for high-resolution imaging-based spatial transcriptomics data and opens new possibilities to study cellular differentiation processes in their spatial context, which helps understanding the natural tissue development.

## Data availability

All datasets used are publicly available data, for convenience preprocessed datasets can be downloaded from Zenodo (https://doi.org/10.5281/zenodo.6798659).

## Code availability

The implementation of SIRV, as well as the code to reproduce the results, are available in the GitHub repository, at https://github.com/tabdelaal/SIRV. The repository source code release is deposited on Zenodo (https://doi.org/10.5281/zenodo.10641057).

## Acknowledgments

This project was supported by the NWO Gravitation project: BRAINSCAPES: A Roadmap from Neurogenetics to Neurobiology (NWO: 024.004.012), Stichting Parkinson Fonds (to R.J.P.), and the NWO TTW project 3DOMICS (NWO: 17126).

