## Supplementary data for "SIRV: Spatial inference of RNA velocity at the single-cell resolution"

Supplementary Data
for

### SIRV: Spatial inference of RNA velocity at the single-cell resolution

Tamim Abdelaal[1,2,3], Laurens M. Grossouw[4], R. Jeroen Pasterkamp[4], Boudewijn P.F. Lelieveldt[1,3], Marcel J.T. Reinders[3,5,6] and Ahmed Mahfouz[3,5,6*]

[1]Department of Radiology, Leiden University Medical Center, 2333ZC Leiden, The Netherlands
[2]Systems and Biomedical Engineering Department, Faculty of Engineering Cairo University, 12613 Giza, Egypt
[3]Delft Bioinformatics Lab, Delft University of Technology, 2628 XE Delft, The Netherlands
[4]Department of Translational Neuroscience, University Medical Center Utrecht Brain Center, Utrecht University, 3584 CX Utrecht, The Netherlands
[5]Department of Human Genetics, Leiden University Medical Center, 2333ZC Leiden, The Netherlands
[6]Leiden Computational Biology Center, Leiden University Medical Center, 2333ZC Leiden, The Netherlands

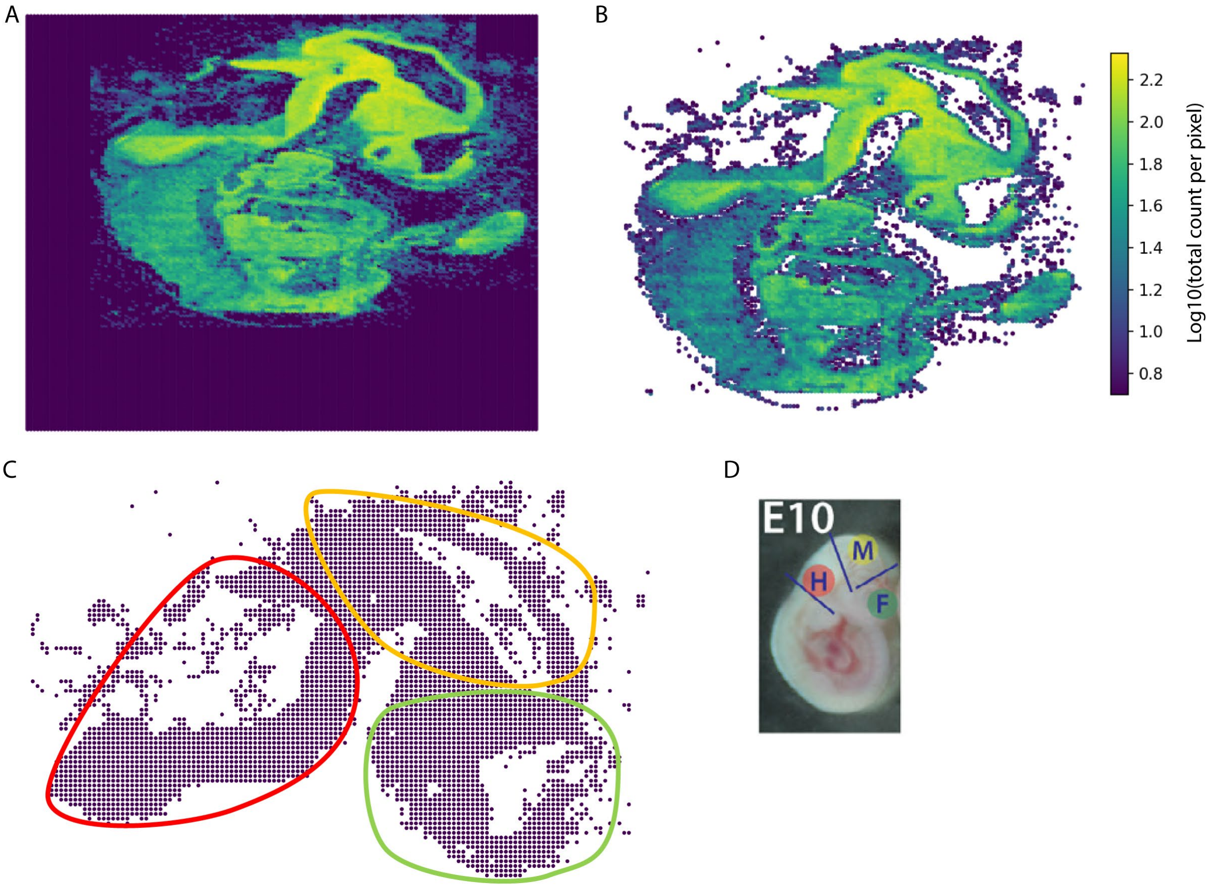


**Supplementary Fig. S1 (A)** Total count per pixel (pseudo-cell) in the HybISS spatial data separating mouse embryonic tissue from background. **(B)** Selecting only tissue pixels with a cutoff of total count per pixel ≥ 4. **(C)** Manual segmentation of only brain tissue (upper part in B), the three brain regions hindbrain (red), midbrain (yellow) and forebrain (green) are highlighted according to (D). **(D)** Figure adapted from G. La Manno et. al[27] illustrating the tissue dissection strategy and highlighting the location of the brain (and different regions in the brain) within the mouse embryo.

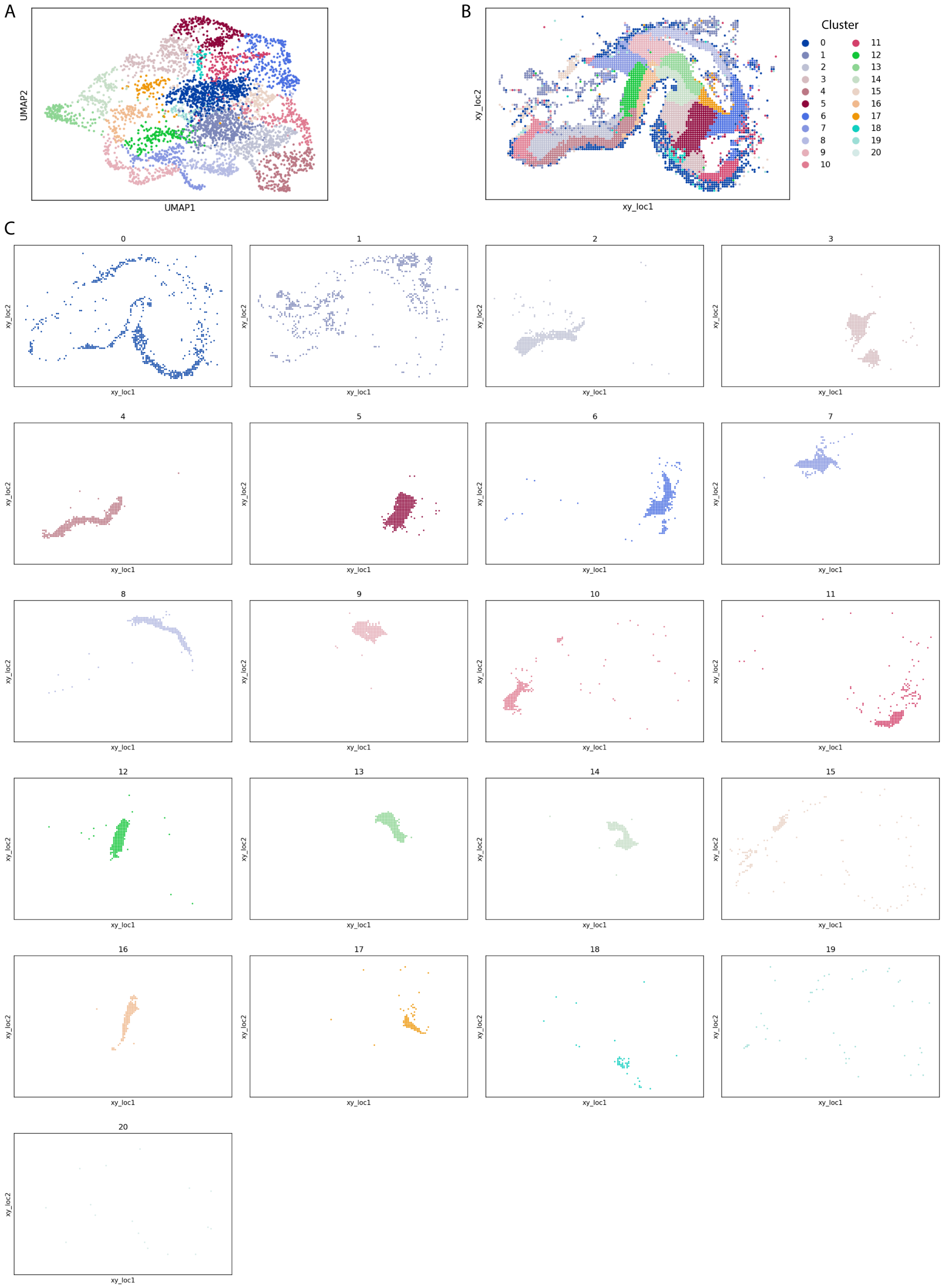


**Supplementary Fig. S2 (A)** UMAP embedding of the HybISS spatial data colored according to 21 cell clusters obtained using Leiden clustering. **(B)** Spatial map of the HybISS data showing spatial localization of the cell clusters. **(C)** Easier visualization of the spatial location of each individual cluster showing one cluster at a time.

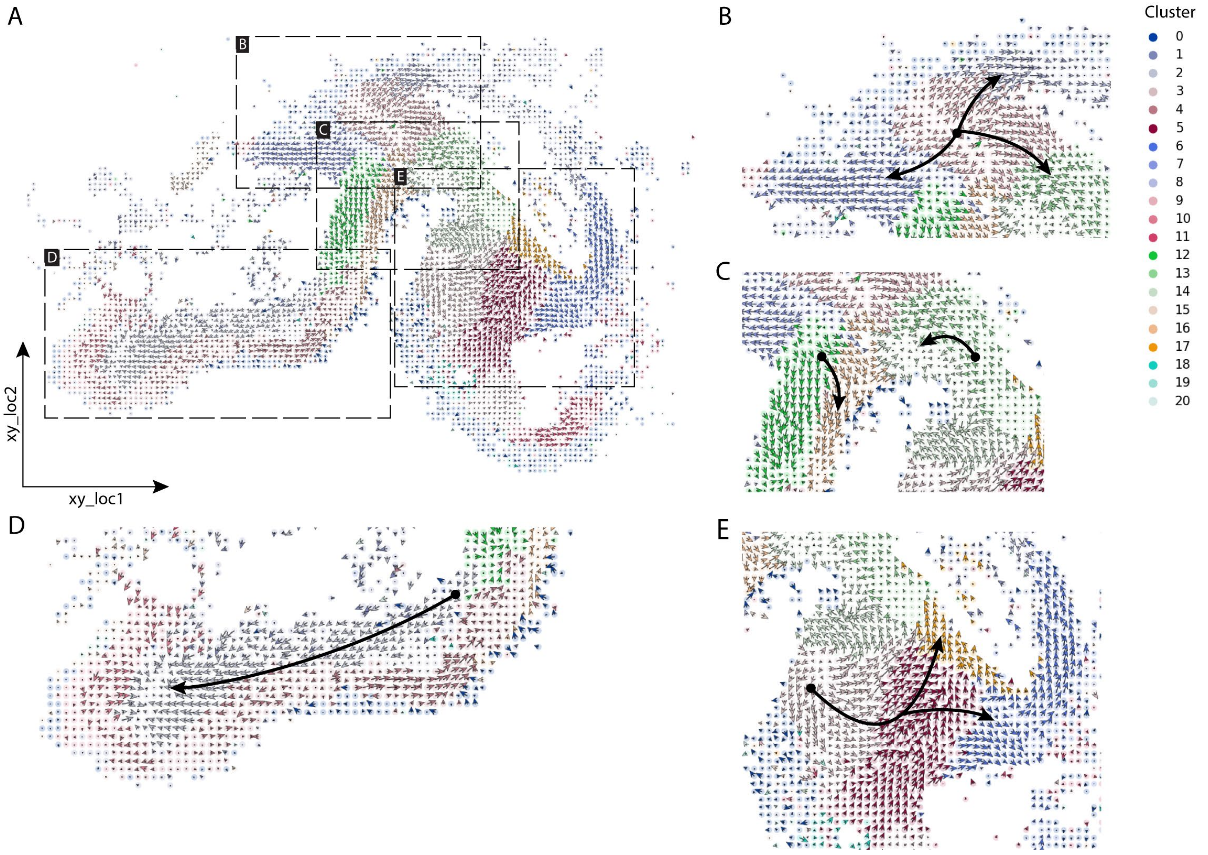


**Supplementary Fig. S3 (A)** Cell-level RNA velocities projected on the spatial coordinates of the HybISS spatial data, colored according to the 21 cell clusters. **(B-E)** Zoom-in views on interesting spatial differentiation trajectories at **(B)** midbrain-hindbrain boundary, **(C)** part of midbrain, **(D)** hindbrain, and **(E)** forebrain. Black arrows show branching or linear spatial differentiation directions between cell clusters (these arrows are drawn manually to highlight certain differentiation trajectories).

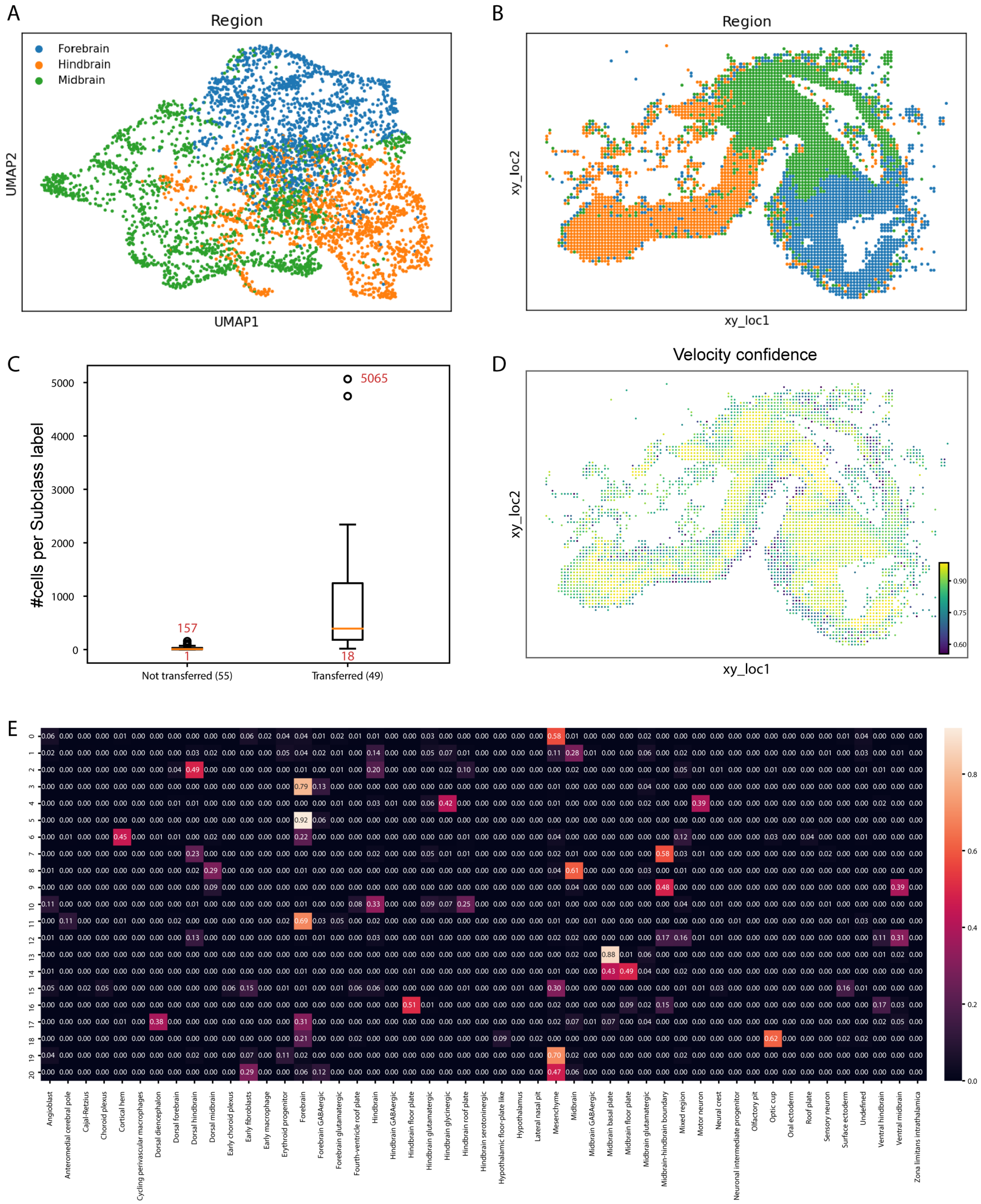


**Supplementary Fig. S4 (A)** UMAP embedding of the HybISS spatial data colored according to the region annotation transferred from the scRNA-seq data. **(B)** Spatial map showing the location of each region label in the tissue. **(C)** Boxplots showing the sizes (in the scRNA-seq data) of the transferred and non-transferred 'Subclass' annotations from the scRNA-seq to the spatial data. **(D)** Velocity confidence score of the obtained spatial RNA velocities using SIRV, visualized over the spatial coordinates of the HybISS spatial data. **(E)** Contingency matrix comparing the HybISS 21 cell clusters (rows) with their corresponding 'Subclass' annotation transferred from the scRNA-seq data (columns). Each row is normalized to sum up to 1.

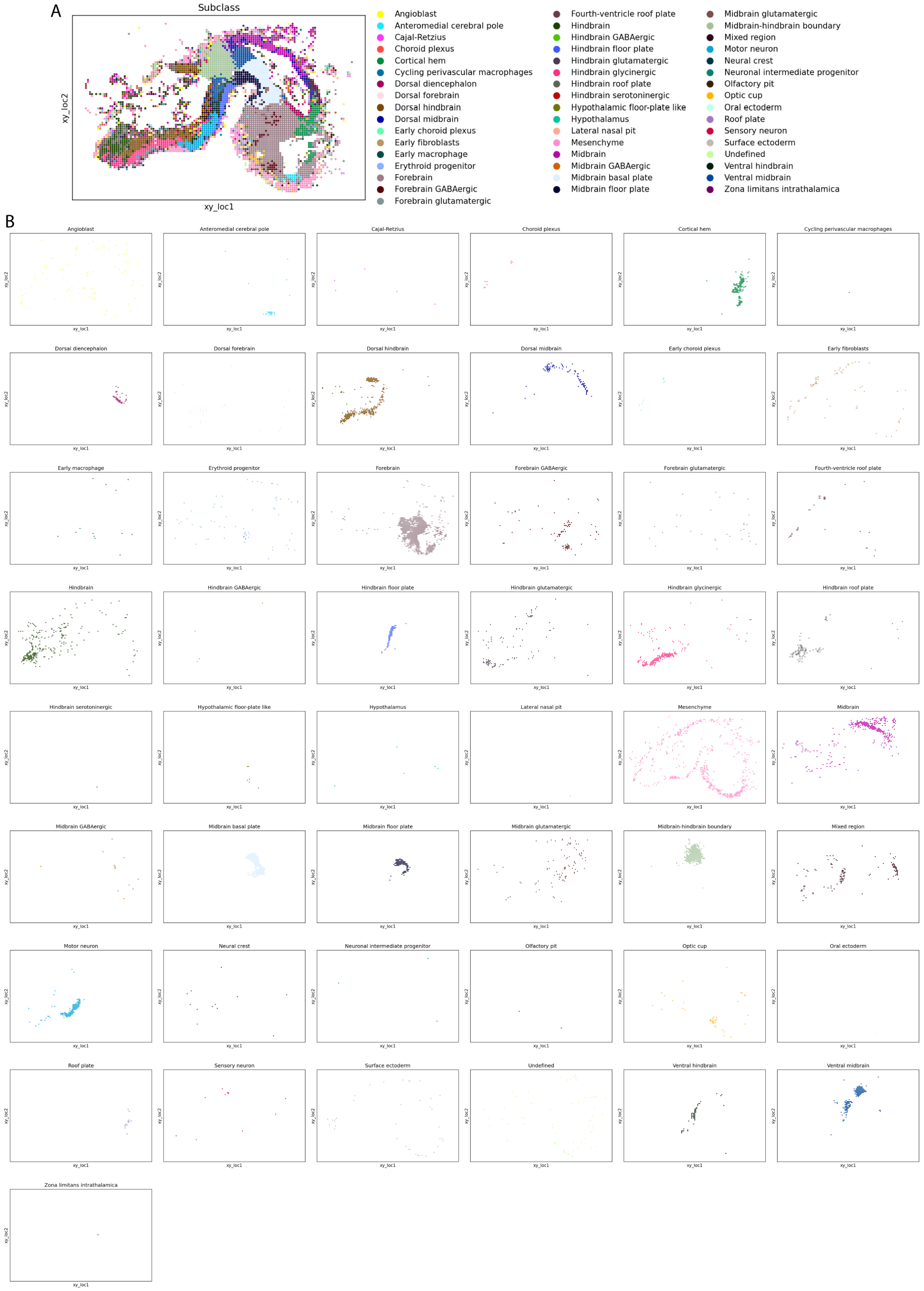


**Supplementary Fig. S5 Spatial distribution of the 'Subclass' annotation transferred from the scRNA-seq data (A)** spatial map showing all 49 subclasses combined, **(B)** clear visualization of the spatial location of each individual subclass showing one subclass at a time.

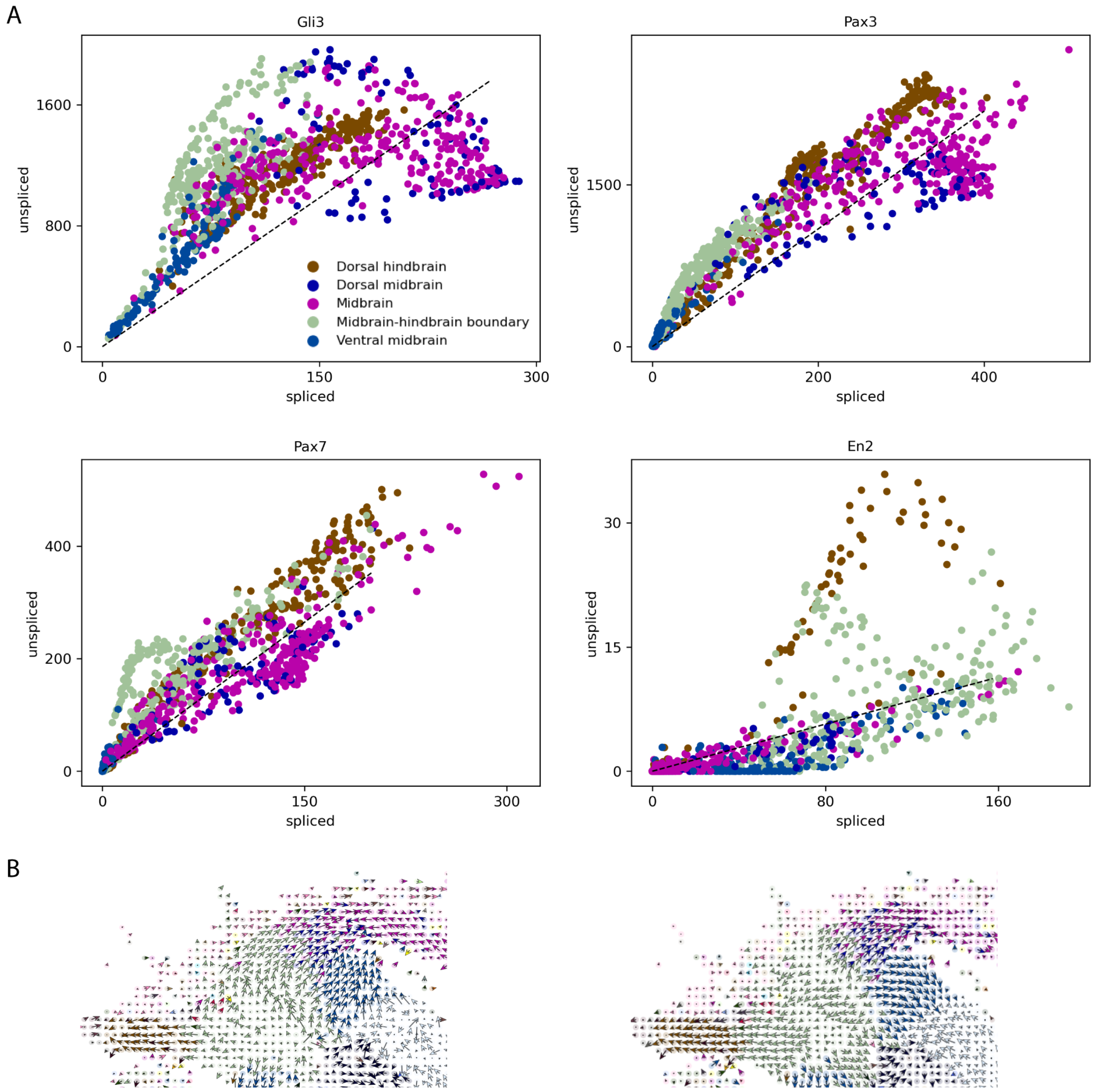


**Supplementary Fig. S6 (A)** Gene velocity plots of *Gli3*, *Pax3*, *Pax7* and *En2* using the subclasses involved in the midbrain-hindbrain boundary differentiation trajectory. **(B)** Cell-level RNA velocities projected on the spatial coordinates of the HybISS spatial data, showing only the midbrain-hindbrain differentiation trajectory. Left plot shows the estimated spatial RNA velocity vectors using only the five genes (*Fgf8*, *Gli3*, *Pax3*, *Pax7* and *En2*) involved into the midbrain-hindbrain differentiation trajectory, right plot shows the estimated spatial RNA velocity vectors using all spatial genes (same plot as Fig. 3B, added again here for easy comparison).

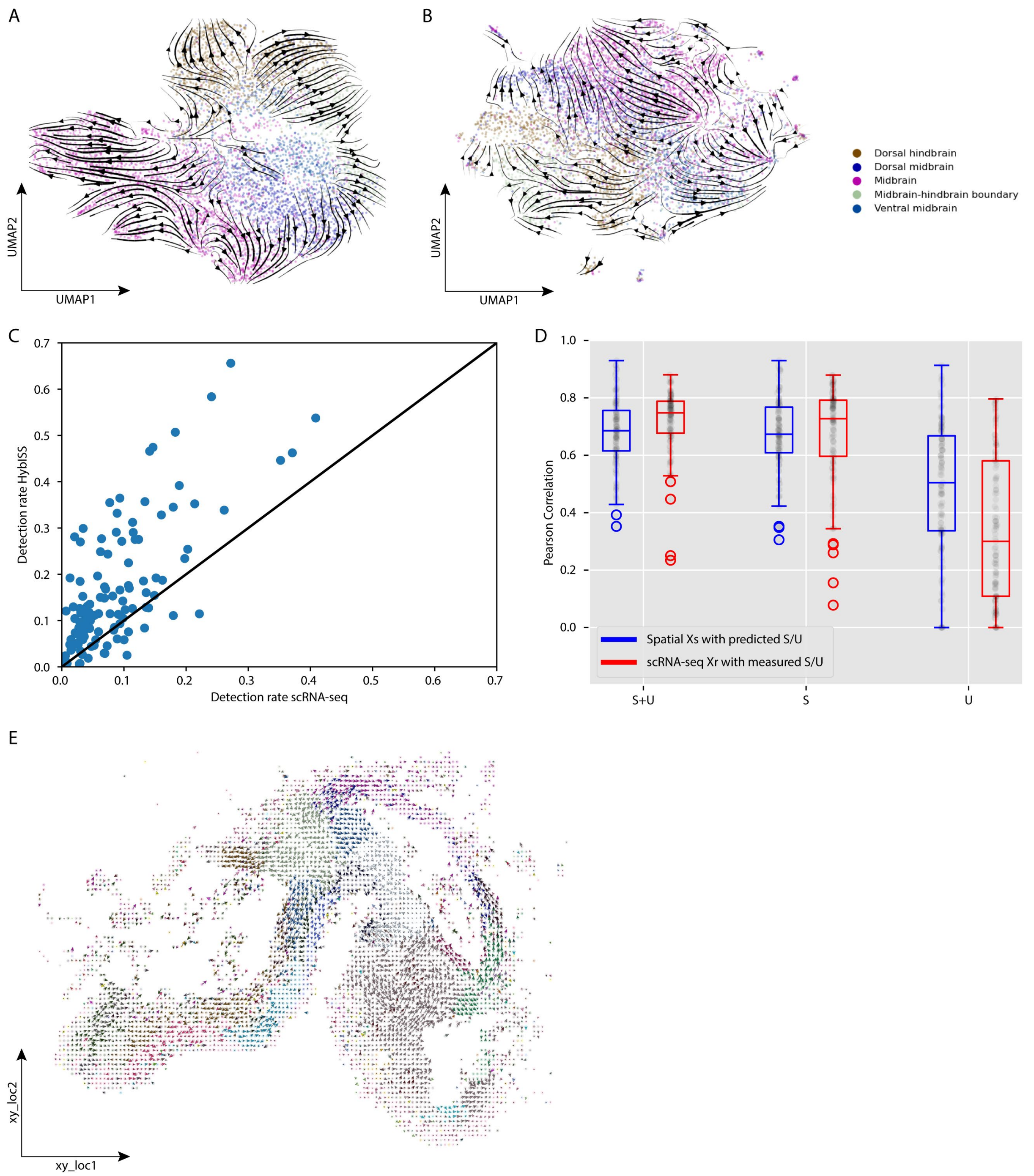


**Supplementary Fig. S7 (A-B)** Main flow of RNA velocities visualized by velocity streamlines projected on the UMAP embedding of a subset from the scRNA-seq data of the developing mouse brain, including the subclasses involved in the midbrain-hindbrain boundary cells differentiation. RNA velocity was calculated using **(A)** top 2,000 HVGs, or **(B)** 117 spatial genes. The branching trajectory of the midbrain-hindbrain boundary cells is not detected correctly in both cases. **(C)** Scatter plot comparing the detection rate of the 117 spatial genes between the HybISS spatial data and the scRNA-seq data of the developing mouse brain. **(D)** Pearson correlation between measured gene expression from the spatial HybISS data and the predicted spliced and unspliced expressions (in blue), which is comparable to correlation obtained between measured gene expression from the scRNA-seq of the developing mouse brain with its measured spliced and unspliced expressions (in red). **(E)** Cell-level RNA velocities projected on the spatial coordinates of the HybISS spatial data, where the measured spatial data was replaced by the summation of the SIRV predicted spliced and unspliced expressions. Cells are colored according to the 'Subclass' annotation transferred from the scRNA-seq data (same legend as Supplementary Fig. S5).

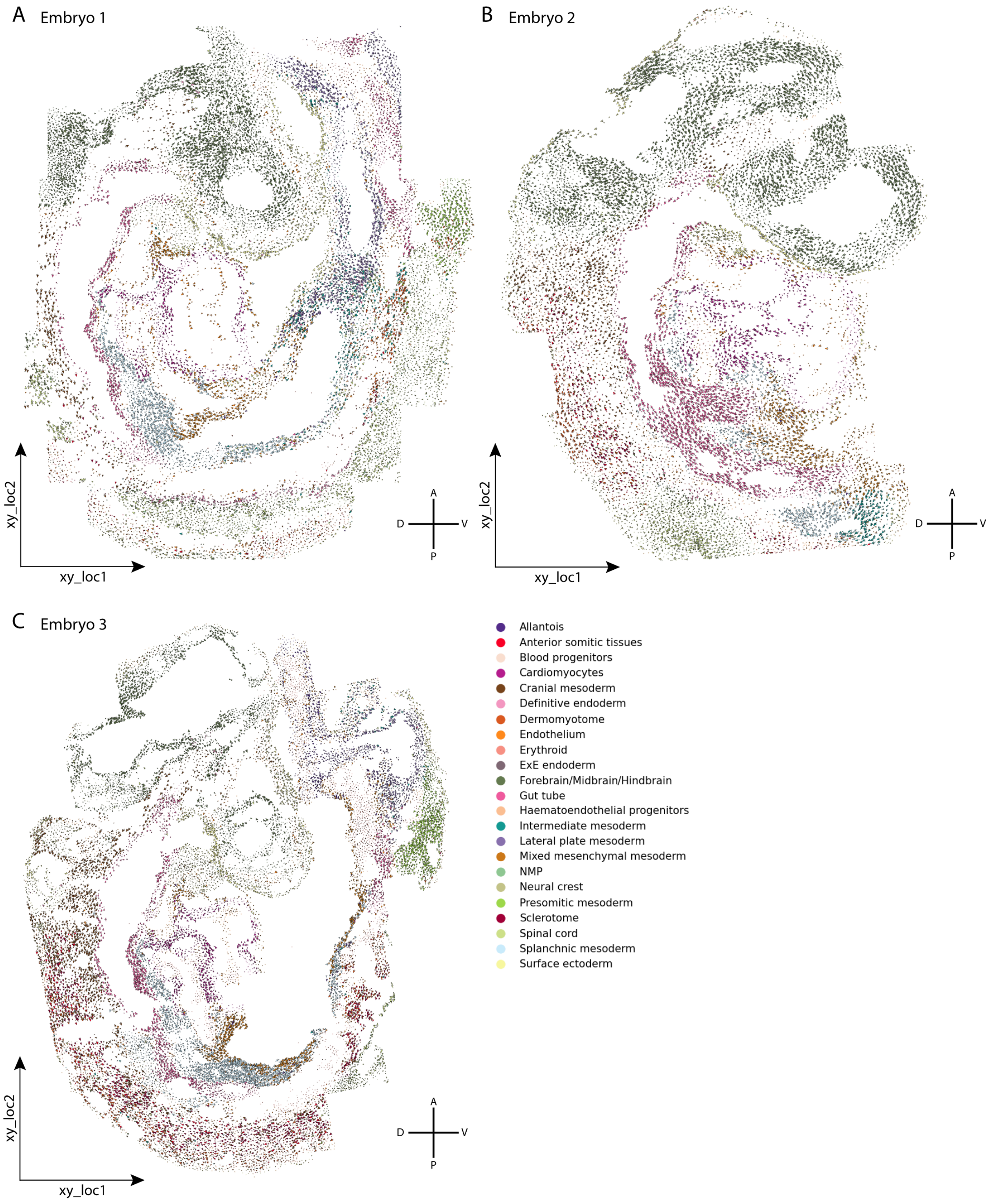


**Supplementary Fig. S8** Cell-level RNA velocities across different cell types projected on the spatial coordinates of the mouse organogenesis SeqFISH data for **(A)** Embryo 1, **(B)** Embryo 2, and **(C)** Embryo 3. Results show reproducible spatial differentiation trajectories across the three embryos.

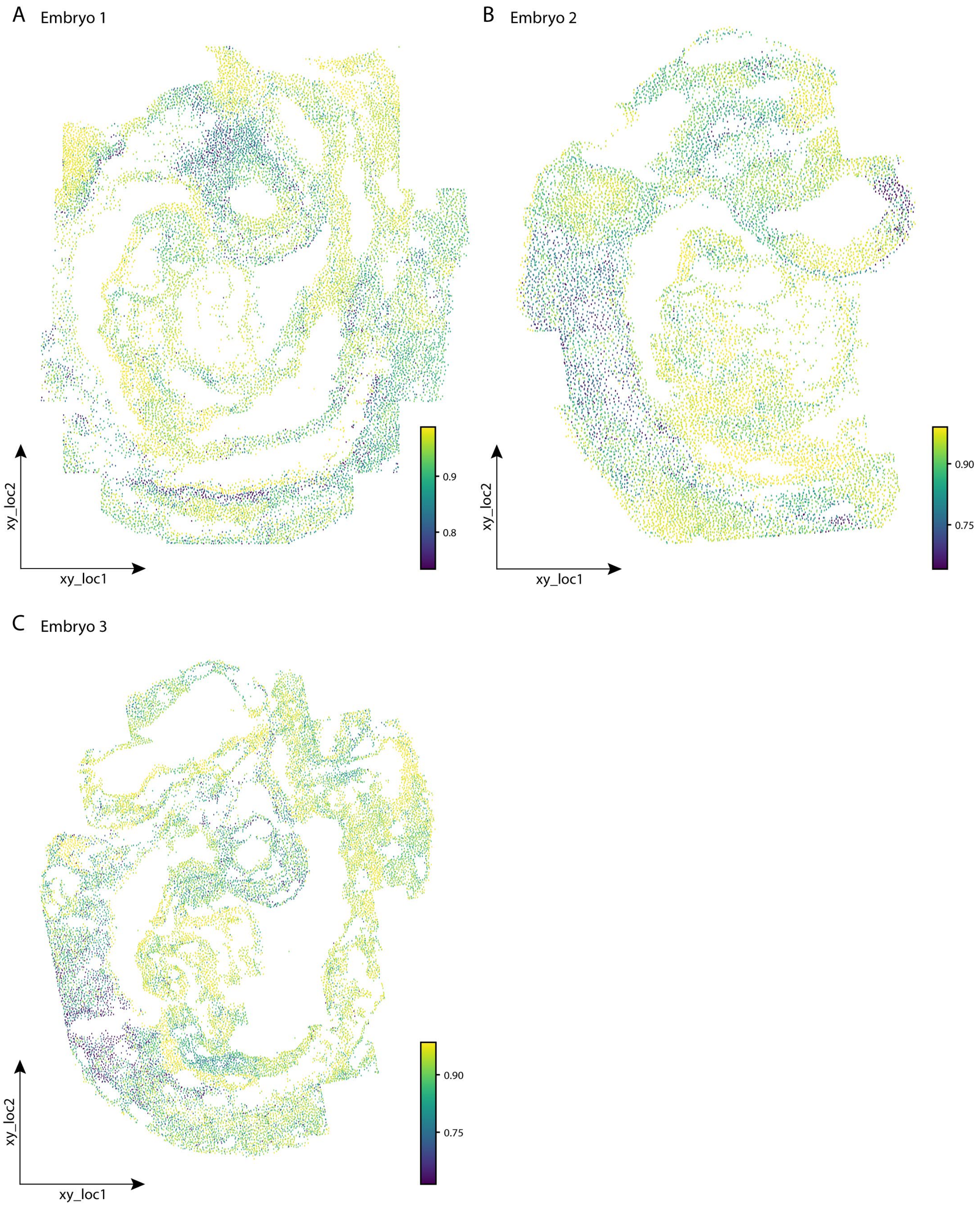


**Supplementary Fig. S9** Velocity confidence score of the obtained spatial RNA velocities using SIRV, visualized over the spatial coordinates of the mouse organogenesis SeqFISH data for **(A)** Embryo 1, **(B)** Embryo 2, and **(C)** Embryo 3.

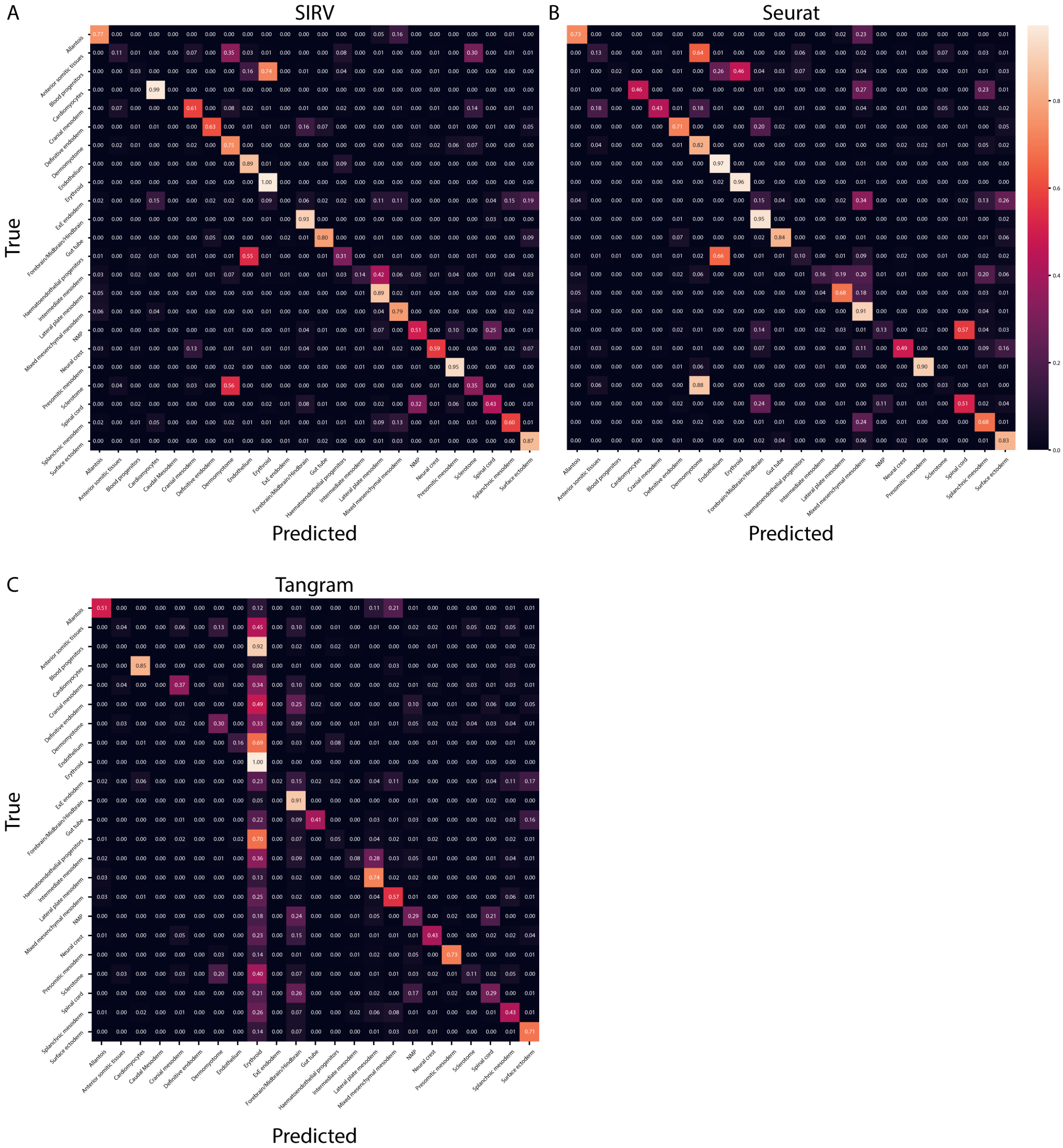


**Supplementary Fig. S10 SIRV label transfer evaluation.** Confusion matrices for **(A)** SIRV, **(B)** Seurat and (C) Tangram. Rows show the true labels, while columns show the predicted labels. Each row is normalized to 1.

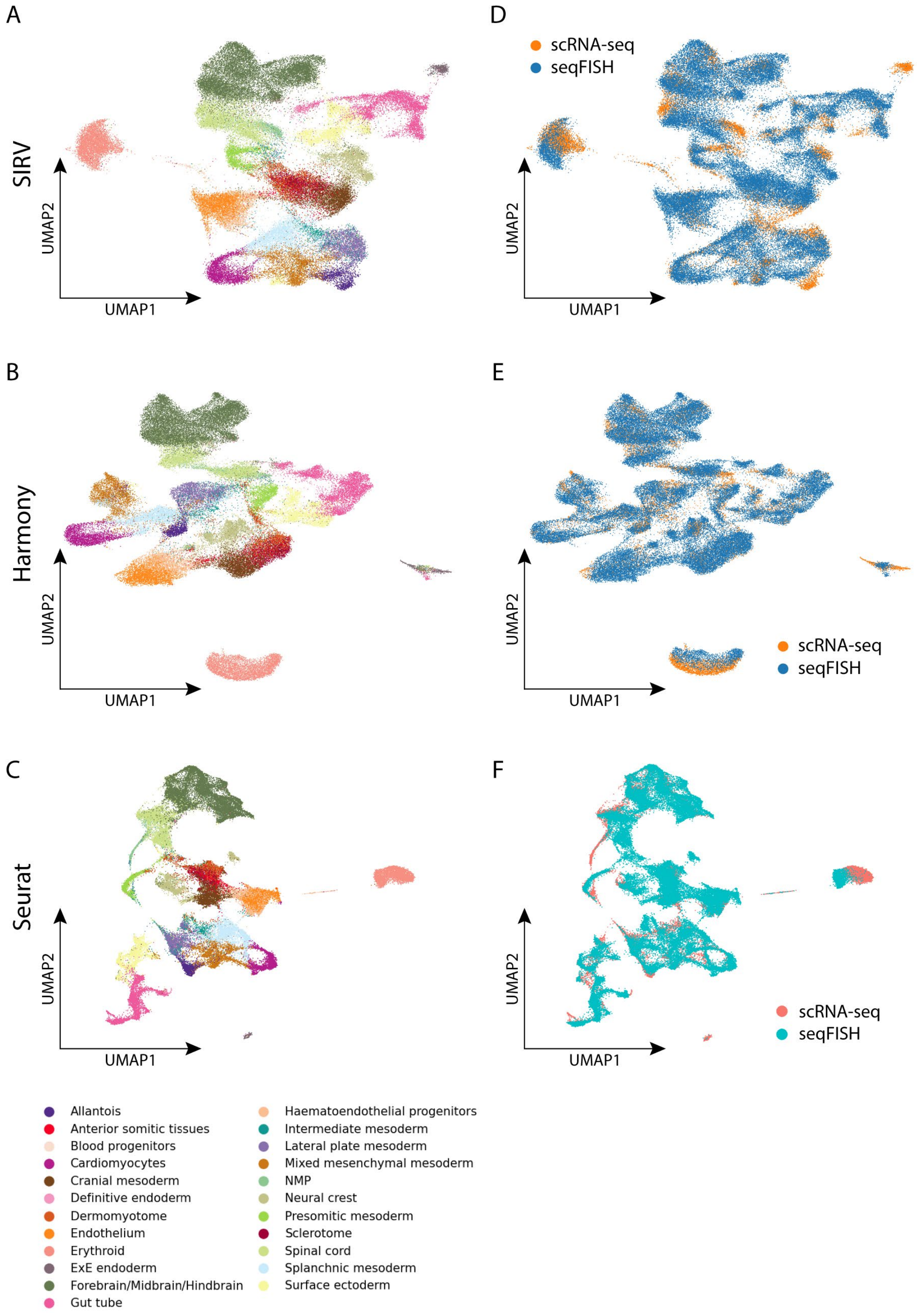


**Supplementary Fig. S11 Qualitative evaluation of the data integration step.** Joint UMAP of the mouse organogenesis Gastrulation scRNA-seq atlas and the seqFISH spatial data using **(A,D)** SIRV, **(B,E)** Harmony and **(C,F)** Seurat, colored by **(A-C)** cell types and **(D-F)** dataset origin.

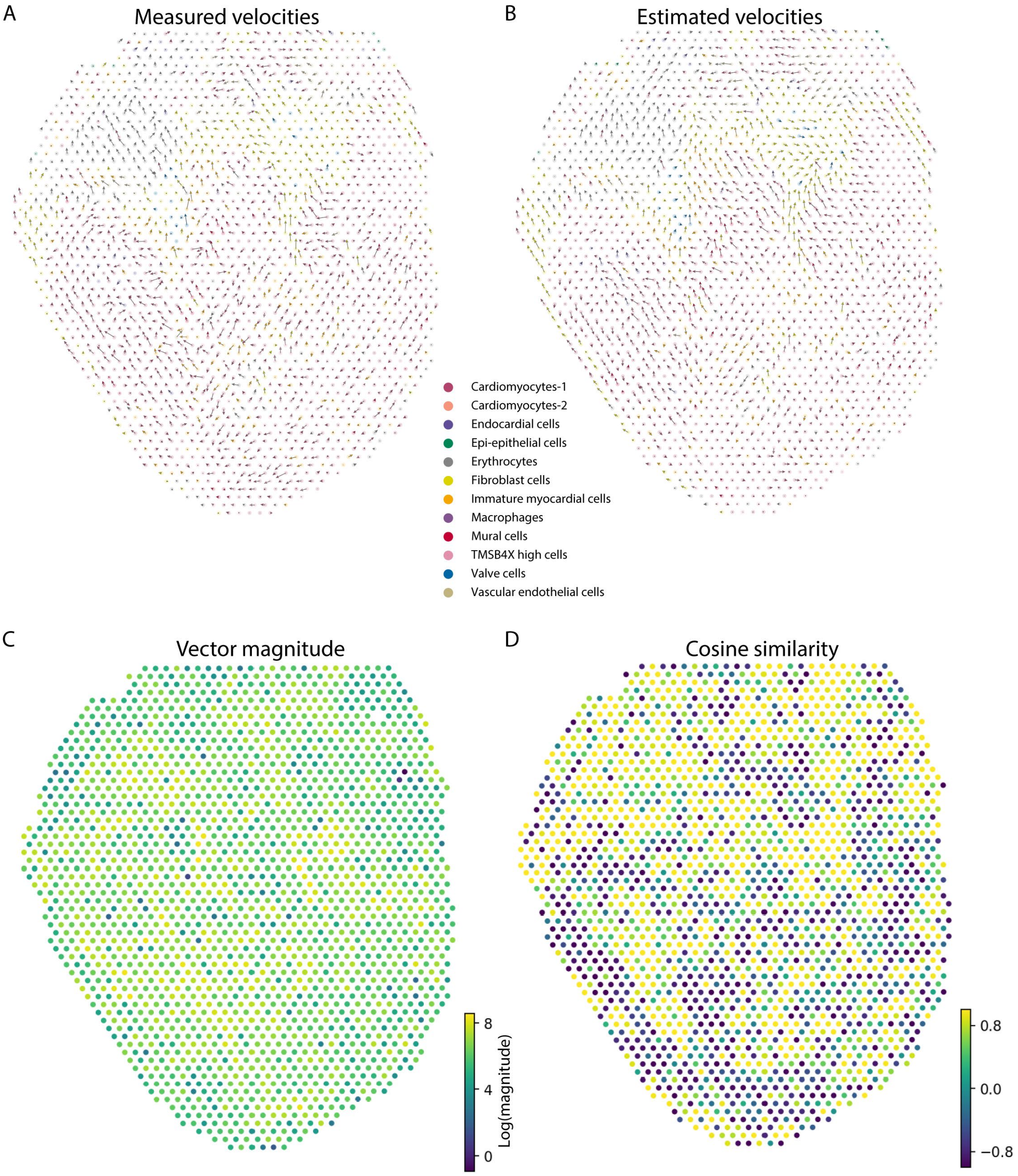


**Supplementary Fig. S12 (A-B)** Spot-level **(A)** measured and **(B)** SIRV estimated RNA velocities across different cell types projected on the spatial coordinates of the developing chicken heart 10X Visium dataset. **(C)** Magnitude of the measured spatial RNA velocity vectors. **(D)** Cosine similarity between measured and estimated spatial RNA velocity vectors at the single-spot level.

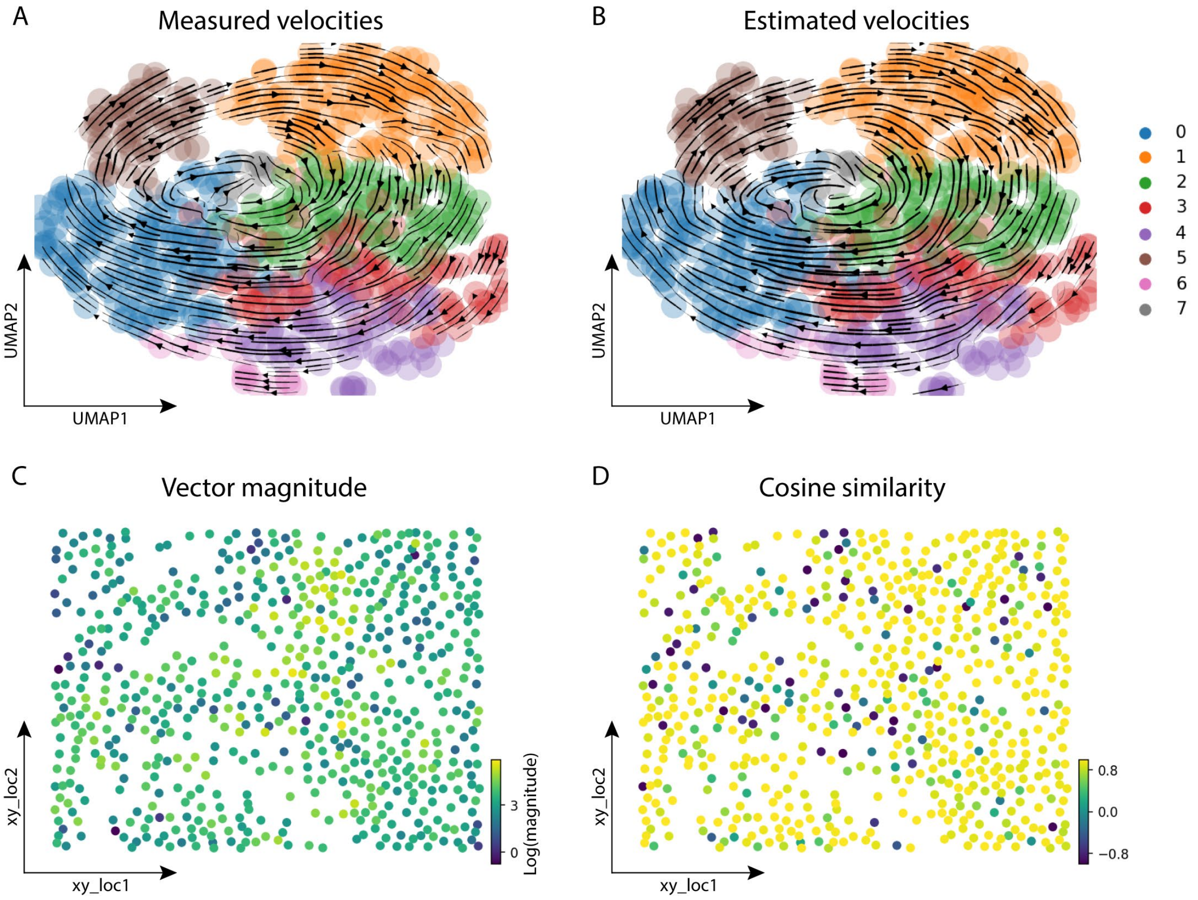


**Supplementary Fig. S13 (A-B)** Main flow of the **(A)** measured and **(B)** SIRV estimated RNA velocities visualized by velocity streamlines, projected on the UMAP embedding of the human osteosarcoma MERFISH dataset, showing high agreement. Cells are colored according to clusters obtained using Leiden clustering. **(C)** Magnitude of the measured spatial RNA velocity vectors. **(D)** Cosine similarity between measured and estimated spatial RNA velocity vectors at the single-cell level.

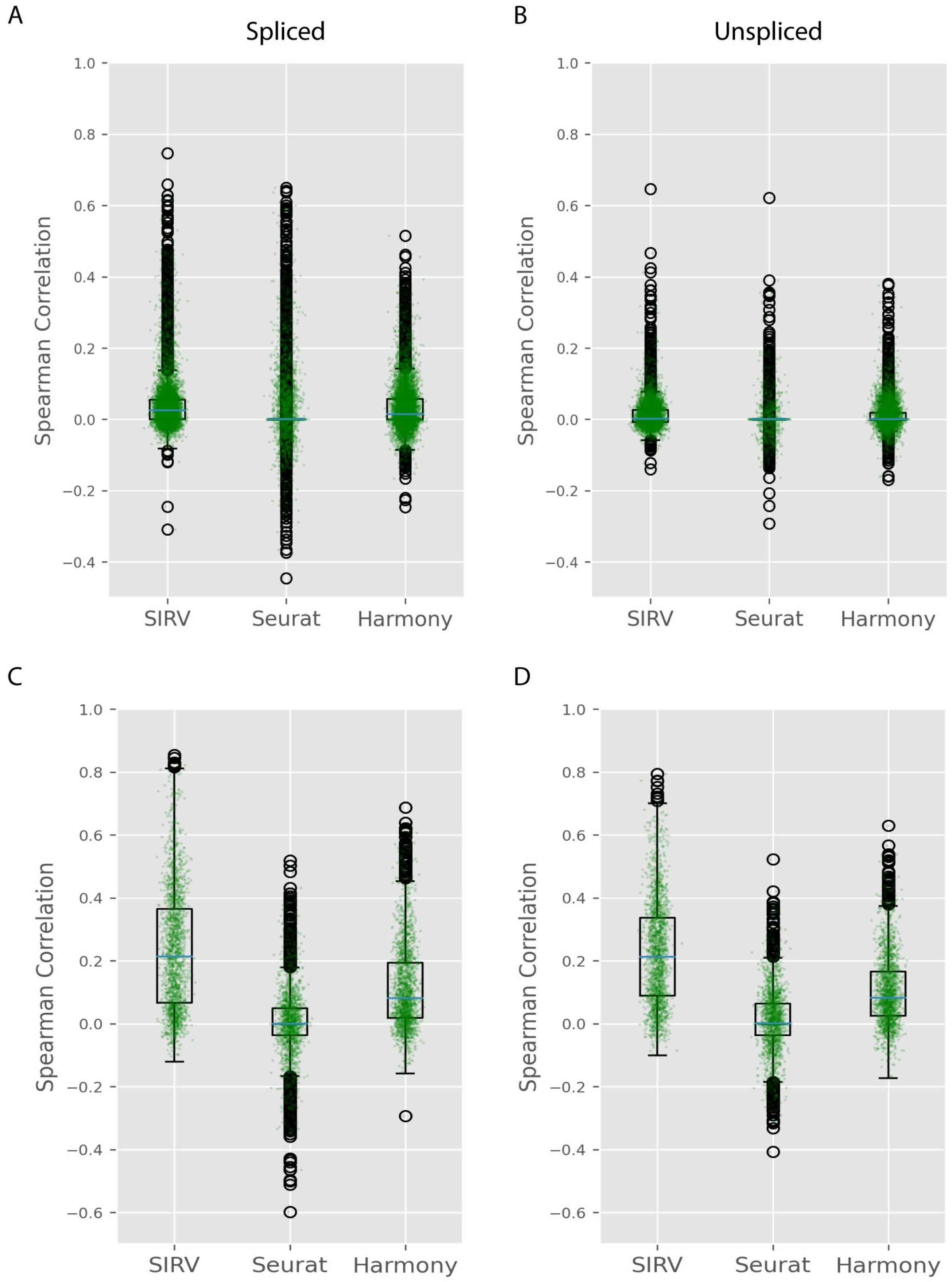


**Supplementary Fig. S14 (A-B)** Boxplots showing the Spearman correlations predicting **(A)** spliced and **(B)** unspliced expressions using SIRV, Harmony and Seurat when applied on the 10x Visium developing chicken heart data. **(C-D)** Boxplots showing the Spearman correlations predicting **(C)** spliced and **(D)** unspliced expressions using SIRV, Harmony and Seurat when applied on the MERFISH data.The blue lines show the median correlation across all genes, and the green dots show the correlation values for individual genes.
